# Global search metaheuristics for neural mass model calibration

**DOI:** 10.1101/2025.08.18.670825

**Authors:** Dominic M Dunstan, Mark P Richardson, Jonathan E Fieldsend, Marc Goodfellow

**Affiliations:** Department of Mathematics and Statistics, University of Exeter, Exeter, UK; Living Systems Institute, University of Exeter, Exeter, UK; Department of Basic and Clinical Neuroscience, Institute of Psychiatry, Psychology, and Neuroscience, King’s College London, London, UK; Department of Computer Science, University of Exeter, Exeter, UK

**Keywords:** neural mass model, EEG, evolutionary search metaheuristics, approximate Bayesian computation, model calibration, parameter inference

## Abstract

Neural mass models (NMMs) are often used to help understand the circuitry that underpins observed brain dynamics in basic and clinical research. A key step is to fuse models with data so that model parameter values can be inferred for a given data set—a process called model fitting or model calibration. This can shed light on putative physiological mechanisms underlying the observed signals.

Calibration is notoriously challenging in biology since models are often non-identifiable, high-dimensional, and nonlinear. Established methods such as dynamic causal modelling (DCM) circumvent some of these issues, for example, by incorporating prior information and employing fast local search methods in the space of feasible parameter values (“parameter space”). However, it is pertinent to better understand the potential limitations of these methods so that we can increase our confidence in the use of models to interpret brain activity, and to develop new approaches as required.

Here we use tools from dynamical systems theory to illustrate some of the complexities of model calibration in an archetypal NMM. We use this information to motivate the use of calibration methods that work across large regions of parameter space, rather than focusing on informative priors or localised search methods. We subsequently evaluate the performance of approximate Bayesian computation (ABC) and evolutionary search metaheuristics (ESMs) for mapping feasible sets of parameters for which an NMM can recreate electroencephalographic recordings during an eyes-closed resting state. Our results demonstrate the superiority of ESMs in terms of computational efficiency and accuracy. Furthermore, we elucidate potential reasons why ESMs are able to perform better than ABC, i.e. that they are less susceptible to biases induced by the complexity of underlying cost landscapes. These results highlight the importance of incorporating ESMs in future efforts to model brain dynamics.

## 1 Introduction

Patterns of brain activity observed in magneto/electro-encephalographic (M/EEG) data are robustly correlated with different cognitive, perceptual or pathological processes occurring in the brain. Observations of brain dynamics can therefore be used to classify brain states or develop biomarkers of disease (Lopes da Silva, 2013). However, observations alone do not help us to understand which underlying neural populations or connectivity patterns are responsible for the dynamics. Combining mathematical models, such as neural mass models (NMMs), with data can help uncover the mechanisms driving observations (Deco et al., 2008). In order to leverage NMMs for the interpretation of M/EEG (and other neuroimaging) data we have to find values of NMM parameters (also referred to as “regions of parameter space”) for which the model can recreate the data. This process is referred to as model fitting or model calibration, and is a crucial component of existing inference and model comparison methods such as dynamic causal modelling (DCM; Kiebel et al., 2008).

Models of biological systems such as the brain present difficult challenges for model calibration, including parameter non-identifiability (multiple parameter values for which the model can accurately recreate data), non-linearity (complex dynamics can emerge with small changes in parameter values), and high dimensionality (Browning et al., 2020; Kirk et al., 2008; Roesch & Stumpf, 2019). The calibration of models of biological systems therefore remains an active field of research that draws on diverse methodology across engineering, mathematics, statistics, and computer science (Gábor & Banga, 2015; Villaverde et al., 2021).

NMMs are often used to understand M/EEG and functional magnetic resonance imaging data, including within the framework of DCM (Friston et al., 2003; Kiebel et al., 2008) and for understanding pathological rhythms in disorders such as epilepsy (Goodfellow et al., 2011; Wendling et al., 2000). NMMs are nonlinear due to the use of a sigmoid function that maps neural population membrane potentials to population firing rates, and as such undergo bifurcations (sudden qualitative changes in model dynamics) as parameters are varied (Cooray et al., 2023; Goodfellow et al., 2011; Grimbert & Faugeras, 2006; Touboul et al., 2011). These bifurcations help to organise the dynamics of the model over changes in parameters, which has implications for model calibration: it means that the regions of parameter space for which the model can recreate the data are likely to be arranged in complicated ways that are difficult to locate. From the perspective of optimisation, to calibrate models, a measure of “fitness”, or inversely “cost”, is required to quantify the difference between model dynamics and data. Bifurcations are known to give rise to cost landscapes that are non-convex and therefore difficult for local optimisation algorithms to navigate (Kirk et al., 2008; Roesch & Stumpf, 2019). Understanding the interplay between bifurcations and cost landscapes is crucial for identifying the limitations of current calibration methods and guiding improvements to ensure that the inferences drawn from mathematical models remain robust.

In this study, we use the Jansen and Rit NMM and power spectral density (PSD) estimated from resting eyes-closed EEG recordings to illustrate the difficulties that arise for commonly used model calibration methods. We draw upon recent evidence to hypothesise that evolutionary search metaheuristics (ESMs) can effectively overcome these issues (Dunstan et al., 2023; Hartoyo et al., 2019; Penas et al., 2024). We benchmark approximate Bayesian computation (ABC) against ESMs using the ground truth cost landscape in parameter spaces of increasing dimensionality, and find that ESMs can overcome sampling biases induced by non-convexities in the cost landscape. We therefore advocate for care when choosing methods to calibrate models of brain dynamics and highlight the potential importance of global search methods like ESMs when prior information on parameter values is unavailable.

## 2 Methods

We focus our analyses on an archetypal NMM developed by Jansen and Rit (Jansen & Rit, 1995; Jansen et al., 1993). For a detailed description of the model, including a model schematic, we refer to Jansen and Rit, 1995. The model and its variants are widely used to simulate EEG activity during various healthy (Jansen & Rit, 1995) and pathological (Wendling et al., 2000) states. Furthermore, it often serves as the foundation of models used in parameter inference methods, including being used as a neuronal model in DCM for EEG (Kiebel et al., 2008).

In short, the Jansen and Rit model is based on the lumped parameter approach developed by Lopes da Silva et al., 1974. It consists of three interacting neural populations: pyramidal neurons, excitatory interneurons and inhibitory interneurons. Activation of each population is described by a second-order differential equation (the pulse-to-wave conversion). Two types of synapses are modelled: excitatory synapses for the pyramidal and excitatory interneuron populations and inhibitory synapses for the inhibitory interneuron population. For *t* ∈ [0, *T*], *T* ∈ ℝ_*>*0_, the model can be described by the following set of coupled differential equations:

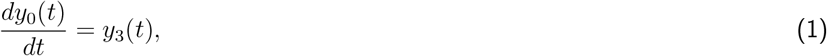

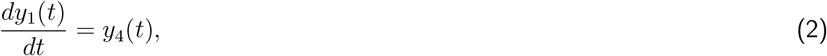

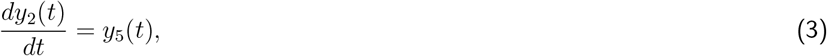

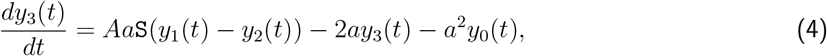

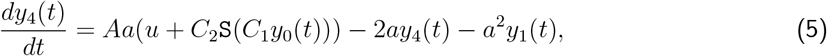

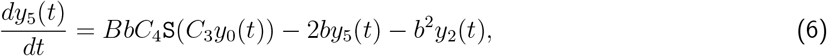

where *y*_0_, *y*_1_, …, *y*_5_ are the state variables of the model. Furthermore, `S`(*v*) = 2*e*_0_*/*(1 + exp(*r*(*v*_0_ − *v*))), *v* ∈ ℝ, converts the mean membrane potential of a population into a mean outgoing firing rate (the wave-to-pulse conversion), and is assumed to act instantaneously. The output of the model is taken to be the membrane potential of the pyramidal population, *y*(*t*) = *y*_1_(*t*) − *y*_2_(*t*). In Equation 5, *u* is an input to the model that represents long-range activity from other unmodelled brain regions, which is often represented by noise. Model parameters, their physiological interpretations, typical values, and the parameter bounds we consider are detailed in Table 1.

**Table 1:** Parameters in the Jansen and Rit NMM. A physiological interpretation, parameter values, and parameter bounds are shown. Individual synaptic connectivity parameters are given by *C*_1_ = *C, C*_2_ = 0.8*C, C*_3_ = 0.25*C* and *C*_4_ = 0.25*C*.

| Parameter | Interpretation | Value | Bounds |
| --- | --- | --- | --- |
| $A$ | Excitatory gain | 3.25 mV | [0 mV, 10 mV] |
| $B$ | Inhibitory gain | 22 mV | [0 mV, 50 mV] |
| $a$ | Excitatory postsynaptic timescale | 100 /s | [25 /s, 140 /s] |
| $b$ | Inhibitory postsynaptic timescale | 50 /s | [6.5 /s, 110 /s] |
| $C$ | Synaptic connectivity | 110 | [0, 500] |
| $v_0$ | Firing rate threshold | 6 mV | [2 mV, 9 mV] |
| $e_0$ | Maximal firing rate | 2.5 /s | [0.5 /s, 7.5 /s] |
| $r$ | Firing rate slope | 0.56 /mV | [0.3 /mV, 0.8 /mV] |
| $u$ | Extrinsic input (mean) | 220 /s | [0 /s, 500 /s] |

In this study, we quantify the model output through the calculation of the power spectral density (PSD) in the model’s linearised form (Moran et al., 2007). This computation is detailed in Sec. 2.2. The model’s PSD is then compared to the PSD estimated from EEG data.

### 2.1 EEG data

We used epochs of 20 s resting eyes-closed EEG data obtained from 10 healthy adults. This data has been used in previous studies (Abela et al., 2019; Dunstan et al., 2023). The recordings were taken on a NicoletOne system at 256 Hz from 19 channels positioned according to the international 10–20 system, with two reference electrodes attached to the ear lobes. Data were re-referenced to the common average. For the purpose of studying the alpha rhythm, we restricted our analyses to the mean of the occipital electrodes. Furthermore, a 2 Hz high-pass *Butterworth filter* was applied to the data to remove low-frequency components. All time series were *z-score* normalised.

### 2.2 Derivation of the Jansen and Rit model transfer function

Here, we detail the derivation of the transfer function of the Jansen and Rit model, enabling the PSD of the model to be analytically calculated from the model parameters. This calculation of the transfer function relies on a linearisation of the model, which indicates that the derivation of the PSD described is only valid in cases where the nonlinear model is well approximated by its linearised form (i.e. in the vicinity of linear dynamical behaviour). To linearise the Jansen and Rit model equations, we restrict attention to fixed points. These fixed points can be found by setting the derivative of state variables in Equations 1 to 6 to zero, obtaining

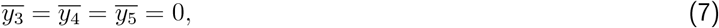

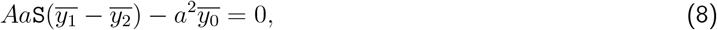

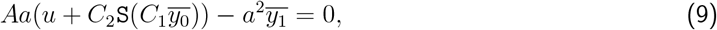

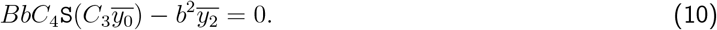

All model parameters are defined in Table 1, 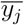 denotes the fixed point value of the state variable *y*_*j*_, for *j* = 0, 1, …, 5, and `S`(*v*) = 2*e*_0_*/*(1 + exp(*r*(*v*_0_ − *v*)), *v* ∈ ℝ. We use Matlab’s `fsolve` function to numerically solve for the values of 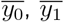 and 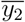. This solver utilises the Trust-Region-Dogleg algorithm (Conn et al., 2000). We generate initial conditions for this solver by simulating the model at a given parameter set for 5 s using Matlab’s `ode45` function, which utilises the Dormand–Prince method (Dormand & Prince, 1980). Zero initial conditions were used to start the model simulation.

In cases where a fixed point has been found, the linearised equations are then obtained by expanding the state variables around their fixed point values. It is convenient to write these equations in state-space form as

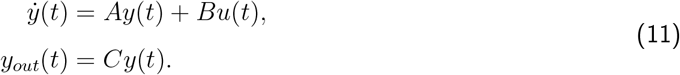

Here, *y*(*t*) and 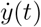 denote the vector of state variables and the derivative vector of state variables, respectively. Furthermore, *A* is the Jacobian matrix of the system of equations evaluated at their fixed point values, *B* is the input mapping vector, denoting how the input to the model (i.e. *u*) is incorporated, *C* is the output mapping vector, denoting that only *y*_1_ and *y*_2_ directly contribute to the model output, which is denoted by *y*_*out*_(*t*). Specifically, these vectors and matrices are defined by the following:

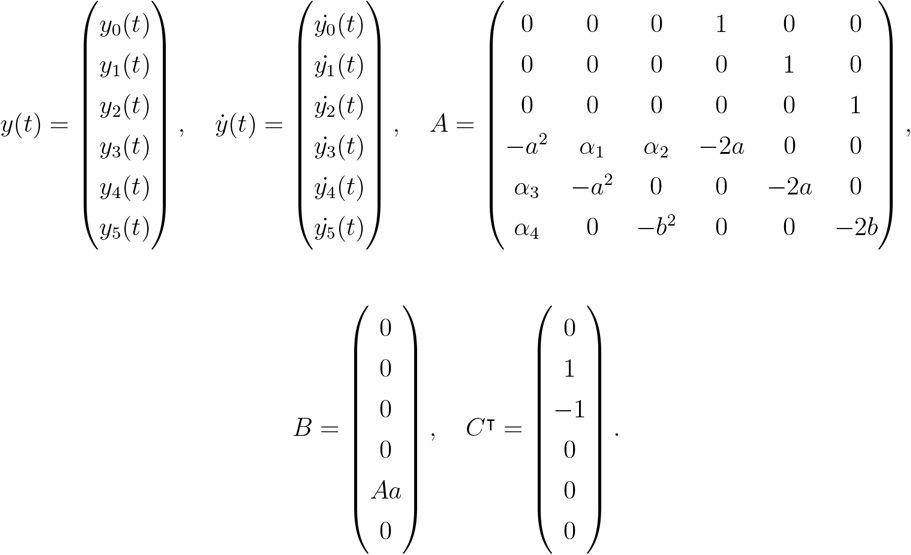

Here, *α*_1_, *α*_2_, *α*_3_, and *α*_4_ are given by,

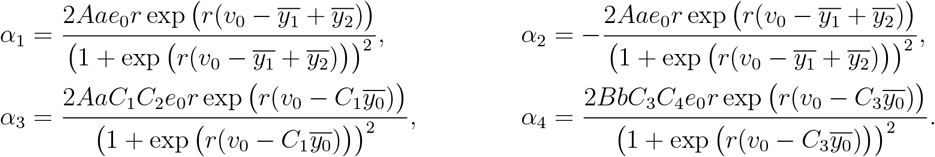

The frequency response and stability of the linearised system can be quantified by taking the Laplace transform of the state space input-output relationship (Moran et al., 2007, 2008; Ogata, 2020). Subsequently, to generate the PSD from the model equations, we take the Laplace transform, ℒ, of Equation 11, with inital conditions *y*(0) = *y*^(0)^, obtaining,

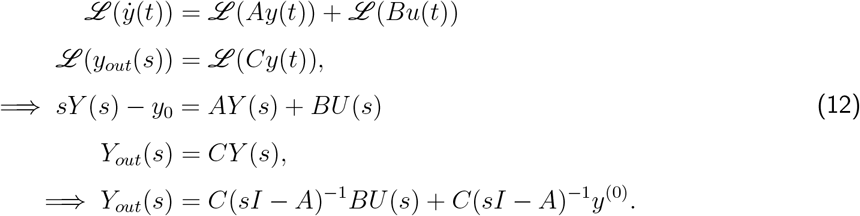

Here, *s* ∈ ℂ, *I* is the identity matrix, and *Y*_*out*_(*s*) is the output of the system in the Laplace domain. Moreover, *Y* (*s*) and *U* (*s*) are the state and input vectors in the Laplace domain, respectively. To generate the spectral response of the system from an input, without loss of generality, we can assume zero initial conditions (i.e. *y*^(0)^ = 0; Ogata, 2020). Moreover, *Y*_*out*_(*s*) is related to the system’s transfer function, *H*(*s*), via *Y*_*out*_(*s*) = *H*(*s*)*U* (*s*). Assuming the spectral response at a stable fixed point, and Gaussian noise input (Wright & Liley, 1994), the spectral response of the model, *S*, at frequency *ω*, can therefore be expressed as,

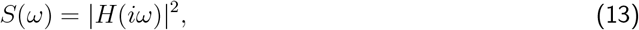

for *H*(*s*) = *C*(*sI* − *A*)^−1^*B* (Ogata, 2020). In this work, we normalise this spectral response to total unit power across a frequency range *ω* = [*ω*_1_, *ω*_2_], thereby obtaining the PSD of the model.

### 2.3 Objective function and tolerance threshold

To compare the dynamics of the model at a given candidate solution to the dynamics observed from the EEG data, we define an objective function (or cost function) as the sum of the squared error between the model PSD (*PSD*_*model*_) and the PSD obtained from the EEG data (*PSD*_*data*_; Dunstan et al., 2023; Hartoyo et al., 2019). Specifically, this objective is defined as,

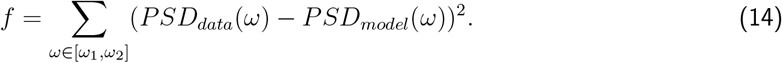

To model the alpha rhythm, which is the dominant rhythm observed on the EEG during an eyes-closed resting state, we select a frequency range of [*ω*_1_, *ω*_2_]=[2 Hz, 20 Hz], with a resolution of 0.0257 Hz. We note that the model and data PSDs were normalised to have total unit power within this range prior to calculating the objective. To compare among algorithms for model calibration, we define a cost tolerance threshold for each subject based on *f*. This enabled us to determine whether a practical alignment between the model and data PSDs had been achieved for a given candidate solution. When a candidate solution had reached this threshold, the algorithm used was terminated, and the parameter set(s) found were retained. We describe this procedure further for each algorithm in Sec. 2.5. Moreover, for each subject, Fig. S1 shows the data PSD and an example model PSD at the defined cost tolerance threshold. To ensure our results were not dependent on the precise threshold value used, we additionally compared the algorithms for a range of cost tolerance thresholds. Figs. S2-S4 show the data PSD and an example model PSD at three other cost threshold values. To model the alpha rhythm, which is the dominant rhythm observed on the EEG during an eyes-closed resting state, we select a frequency range of [*ω*_1_, *ω*_2_]=[2 Hz, 20 Hz], with a resolution of 0.0257 Hz. We note that the model and data PSDs were normalised to have total unit power within this range prior to calculating the objective. To compare among algorithms for model calibration, we define a cost tolerance threshold for each subject based on *f*. This enabled us to determine whether a practical alignment between the model and data PSDs had been achieved for a given candidate solution. When a candidate solution had reached this threshold, the algorithm used was terminated, and the parameter set(s) found were retained. We describe this procedure further for each algorithm in Sec. 2.5. Moreover, for each subject, Fig. S1 shows the data PSD and an example model PSD at the defined cost tolerance threshold. To ensure our results were not dependent on the precise threshold value used, we additionally compared the algorithms for a range of cost tolerance thresholds. Figs. S2-S4 show the data PSD and an example model PSD at three other cost tolerance threshold values.

We calculated the model’s PSD directly from the linearised equations (as described). For the EEG data, the PSD was estimated using Welch’s method (Welch, 1967). To implement this, we split each 20 s data epoch into 8 segments with 50% overlap. We note that, compared to simulating the model time series and estimating the PSD using Welch’s method, a vast computational speed-up is achieved by calculating the PSD of the model directly from its linearised form. This approach comes with the caveat that the approximation is not valid for nonlinear dynamical behaviour. Many studies commonly analyse the linear dynamical properties of NMMs (Hartoyo et al., 2019, 2020; Moran et al., 2007). For example, DCM often employs spectral features to compare model dynamics with data (Kiebel et al., 2008; Moran et al., 2007, 2008). In this study, we adopt this approach to characterise the dynamics, as it provides computational tractability for a detailed comparison of the algorithms we use.

### 2.4 Bifurcation analysis

Prior to comparing algorithms to map model parameters from data, we performed a bifurcation analysis of the Jansen and Rit model to investigate the qualitative changes in the dynamics of the system as model parameters are varied. We note that we focused on local bifurcations and that the bifurcation analysis was conducted in the noise-free ODE version of the Jansen and Rit model (i.e. *u*=constant in Equation 5).

We identify three types of bifurcations in our analysis: saddle-node bifurcations, Hopf bifurcations, and Bogdanov-Takens bifurcations. Briefly, a saddle-node bifurcation occurs when a stable and unstable fixed point coalesce and annihilate each other as a parameter is varied. A Hopf bifurcation occurs when a pair of complex conjugate eigenvalues of the Jacobian matrix cross the imaginary axis. A Hopf bifurcation is called a supercritical Hopf if a stable fixed point loses stability and generates a stable limit cycle (or another stable attractor; Izhikevich, 2007). Both saddle-node and Hopf bifurcations have codimension 1, meaning they can occur by varying a single model parameter. Conversely, a Bogdanov-Takens bifurcation has codimension 2. This bifurcation is characterised by the simultaneous occurrence of a saddle-node and Hopf bifurcation, and occurs when the Jacobian matrix has two zero eigenvalues at the bifurcation point (Izhikevich, 2007). We used Matcont (Dhooge et al., 2003) to conduct the bifurcation analysis presented herein and numerically track bifurcations from varying parameter *A* (1-dimensional bifurcation) and parameters *A* and *a* (2-dimensional bifurcation) in the model.

### 2.5 Methods for model calibration

In total, we used four different approaches to find the regions of parameter space for which the simulated PSD sufficiently models the PSD observed in the EEG data. The algorithms we used were ABC, a genetic algorithm (GA) and the niching migratory multi-swarm optimiser (NMMSO), along with a Latin hypercube sampling (LHS) of the parameter space. We implemented these algorithms in three scenarios, varying 2, 3 and 9 parameters in the Jansen and Rit model. Here we outline each of these algorithms.

ABC is a Bayesian method developed for statistical inference. ABC uses simulation-based inference, which bypasses the evaluation of the likelihood function (Sunnåker et al., 2013). This is especially advantageous in complex models when the likelihood function is difficult (or potentially impossible) to analytically define. Here, we used a common variant of ABC called sequential-Monte Carlo ABC (ABC-SMC; Toni et al., 2009). In general, this method involves treating the parameters of the model as random variables whose probability distributions are unknown. The algorithm then aims to recover the posterior distribution of the parameters, given the data and some prior distribution. The likelihood function is replaced by forward simulations of the model and a comparison between the observed and simulated data. This comparison is made by defining a summary statistic. In this study, we use uninformative priors by setting the prior distribution for each parameter that was varied to a uniform distribution between its box constraints (West et al., 2021; see Table 1 for a list of bounds used for each parameter). The algorithm proceeds by randomly taking a sample of draws from the prior distribution and iteratively accepting or rejecting a simulation of the forward model (the calculation of the model PSD at a given parameter set). The acceptance or rejection criterion is based on defining a distance measure from the summary statistic to a predefined tolerance value. To compare ABC directly with the other algorithms, we defined the summary statistic as the computed PSD and the distance measure as how accurately the model PSD reproduces the data PSD (i.e. as defined by Equation 14). In this study, we used a shrinking tolerance schedule of tol = [5*f*_1_,2*f*_1_,1.5*f*_1_,*f*_1_], for *f*_1_ the final tolerance threshold value. We note that Fig. S1 defines the final tolerance threshold used for each subject, with Figs. S2–S4 showing additional tolerance threshold values that were tested. Bayes’ Theorem is applied for each iteration of this tolerance. We used a Gaussian permutation kernel to generate the particles for each ABC-SMC generation (Toni et al., 2009). The shrinking tolerance applied enables a higher acceptance rate for the algorithm. It is important to highlight that any tolerance schedule involves a trade-off between computational cost and accuracy. Furthermore, following previous studies of ABC in neural mass modelling (West et al., 2021), we used a particle size of *N* =512. Therefore, when *N* =512 points have been found from the final tolerance, the algorithm is terminated and the distribution of parameters that simulated the data (the posterior) is retained. We refer to Toni et al., 2009 for a detailed description of ABC-SMC. Furthermore, we refer to West et al., 2021 and Ditlevsen et al., 2023 for ABC-SMC applications specific to neural mass modelling from electrophysiological data. Here we used the Julia toolbox *GpABC* (Tankhilevich et al., 2020) for a fast implementation of the ABC-SMC algorithm.

In addition to ABC, we also used two ESMs to infer the model parameters that recreate the PSD observed in the data. Specifically, we used a GA (Michalewicz, 1996) and the NMMSO method (Fieldsend, 2014). Both these algorithms are single-objective. We implemented these algorithms to minimise the objective function defined in Equation 14. We used box constraints in the decision space based on minimum and maximum feasible parameter values (as defined by the bounds in Table 1).

For the GA, we chose to use a population size of 100, 150 and 500 individuals across 2, 3 and 9 model parameters, respectively. Each individual in the population was evaluated using the cost function defined by the objective in Equation 14. The population at the initial generation was obtained through an LHS of the parameter space. To obtain the population of the subsequent generations, we used Matlab’s standardised operations in the function `ga` for elitism, mutation and crossover. We set the algorithm to iterate through generations until a candidate solution was obtained that provided a cost value below the threshold defined for the given subject (denoted *f*_1_). When this occurred, we terminated the algorithm and retained the corresponding parameter set. To compare the inference results with other algorithms, for each optimisation problem, we ran *N* =512 separate GAs to obtain a distribution of parameters that describes the data.

Along with GAs, other ESMs have been used for recovering NMM parameters from data. This includes particle swarm optimisation (PSO; Kennedy and Eberhart, 1995), which has shown promise in recovering parameters from EEG (Hartoyo et al., 2019, 2020; Shan et al., 2016). Here, we used a multi-swarm variant of a PSO algorithm called NMMSO (Fieldsend, 2014). This algorithm is a *multi-modal* metaheuristic, designed to explore the decision space to find all candidate solutions that locally minimise a given objective. The algorithm implements this approach by developing sub-swarms that search across separate modes in the decision space simultaneously, and returns a set of estimated local optima locations. We refer to Fieldsend, 2014 for further information on the NMMSO. Here, we implemented the algorithm to find parameter sets that simulate the PSD observed in the data. We defined the swarm size (i.e. the maximum number of particles per swarm) as *λ* = 4 + ⌈3*log*(*d*)⌉, for *d* the dimension, or equivalently, the number of parameters being varied (2, 3 or 9; following the population scaling approach of Hansen, 2006). Similarly to the GA, we set the algorithm to iterate particle positions until we found (at least) one solution with a cost below the defined threshold (*f*_1_) for the given subject. We repeated this approach until we obtained *N* =512 solutions that formed a distribution of parameters that describe the data.

Finally, we conducted an LHS of the model parameters. This method ensures that each input variable is sampled uniformly across its entire range while aiming to achieve an efficient space-filling design. We simulated an LHS with a size of 1.5 million, 4.2 million, and 25 million points when varying 2, 3 and 9 model parameters, respectively. This sampling size was set large enough to ensure *N* =512 points under the defined cost tolerance threshold (*f*_1_) were obtained. Unlike the previous algorithms, this method does not ‘search’ the landscape to find solutions which can recover parameter sets that simulate data. Alternatively, this approach provided an unbiased estimate of the ‘true’ parameter distributions that can recreate the data (i.e. an estimate of the ground truth). We therefore used the LHS as a benchmark to test how accurately each algorithm performed the parameter inference on the EEG data. We note that, for each algorithm used, we additionally retained and compared the number of function evaluations required to obtain the parameter distributions.

### 2.6 Jensen-Shannon divergence

The Jensen-Shannon divergence (JSD) is a measure of similarity between two probability distributions.

It is a symmetrised and smoothed version of the Kullback-Leibler divergence (Nielsen, 2019). Given probability distributions *P* and *Q* in domain Ω and Shannon entropy *H*(*P*) = − ∑_*x*∈Ω_ *P* (*x*)*log* (*P* (*x*)), for *x* ∈ ℝ, then,

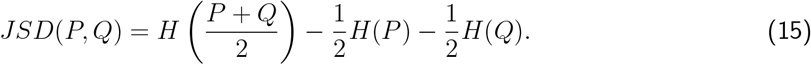

In this study, for each algorithm and each subject, we recovered parameter distributions comprising *N* =512 values. We estimated probabilities from these values using 100 equally sized bins. For each parameter, we then applied the JSD as a measure for the similarity of the distribution obtained between the algorithm we test (i.e. ABC, GA or NMMSO), and our unbiased estimate of the ‘true’ distribution the parameter can take, given the data (which is found through the LHS). Consequently, for a parameter distribution obtained from a given algorithm, lower values of the JSD indicate a more accurate estimation.

### 2.7 Test cost function

Finally, in addition to the analysis conducted using the Jansen and Rit model, we tested the parameter inference algorithms on a simple 1-dimensional cost function. We designed this cost function to test how well the algorithms deal with suboptimal solutions in the cost landscape, and to provide further validation of the differences between the algorithms found. For a single input variable, *x*, this function was defined as, *h*(*x*) = |*sin*(15*x*) + *sin*(12*x*) + *sin*(10*x*)|. We set the bounded box constraint on *x* to be *x* = [0.6, 1.4]. Furthermore, based on visual inspection, we defined a cost threshold to be *f*_2_=0.08, which determined when the algorithms were terminated. To implement ABC, we iterated through a shrinking tolerance schedule of tol = [5*f*_2_,2*f*_2_,1.5*f*_2_,*f*_2_]. In general, for each algorithm, we used the same approaches as described previously, but with an LHS of 20000 points and a population size of 16 individuals in the GA. Moreover, to avoid any potential small sample biases, we implemented 10 repeats of each algorithm on this function and retained the mean values from these repeats.

## 3 Results

### 3.1 Illustrating non-convexities in neural mass model cost landscapes

The output of nonlinear dynamical systems is organised by invariant sets and bifurcations. To illustrate this for NMMs, Fig. 1A shows a bifurcation diagram of the Jansen and Rit model as parameter *A* is varied. The bifurcation diagram shows 2 supercritical Hopf bifurcations that generate a stable limit cycle between approximately *A*=4.55 and *A*=9.35, along with 2 saddle-node bifurcations at approximately *A*=2.30 and *A*=2.42. The presence of Hopf and saddle-node bifurcations is expected in nonlinear NMMs and has been observed previously (Goodfellow et al., 2011; Spiegler et al., 2010; Touboul et al., 2011), including in the Jansen and Rit model (Grimbert & Faugeras, 2006). To illustrate how these bifurcations organise model dynamics, we visualised the model’s PSD for *A*=4.25 and *A*=9.52, which are on either side of the supercritical Hopf bifurcations and are shown in Fig. 1B and Fig. 1C, respectively. Note, these values are also displayed by vertical dotted lines in Fig. 1A and Fig. 1D. It can be seen that the PSDs have a very similar shape for these different values of parameter *A*, with both PSDs generating alpha rhythm dynamics. We superimposed the PSD from an example subject (subject 1) onto the model PSDs in Fig. 1B and Fig. 1C. The comparison shows that the PSDs generated by the model were visually similar to the PSD estimated from this subject’s EEG data.

**Figure 1:**
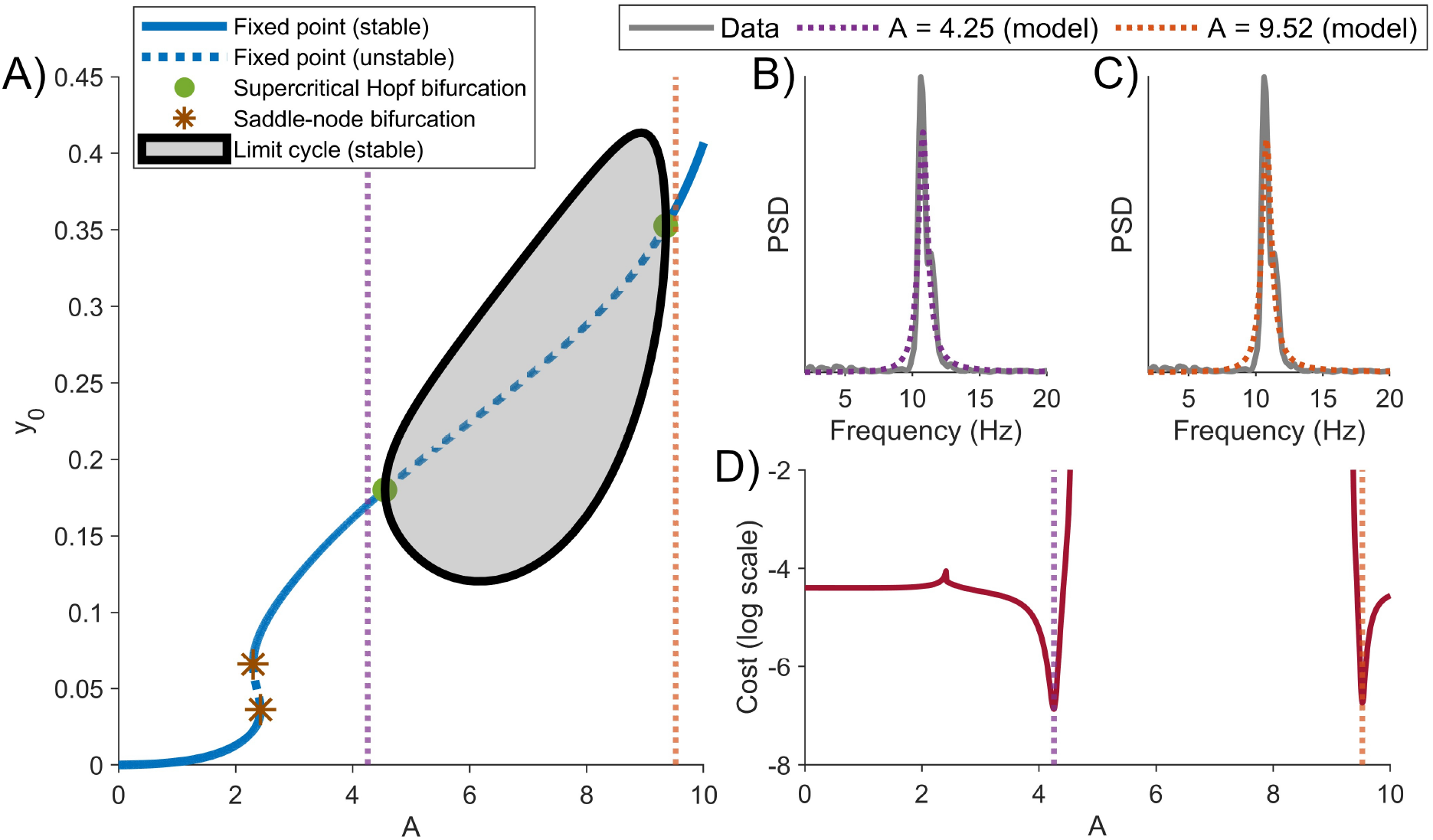
Bifurcation analysis and cost landscape of the Jansen and Rit model as parameter A is varied. A) Bifurcation analysis of the Jansen and Rit model as parameter *A* is varied. Other model parameters were set to their typical values, as defined in Table 1. The bifurcation diagram is shown for state variable output *y*_0_. Note that this state variable was used for simplicity, but the model output is *y* = *y*_1_ − *y*_2_. 2 saddle-node bifurcations and 2 supercritical Hopf bifurcations occurred across this range of parameter *A* (see legend). The supercritical Hopf bifurcations produce a stable limit cycle between *A*=4.55 and *A*=9.35. The minimum and maximum amplitude of this stable limit cycle is indicated by the black line. The PSD from subject 1 and the model PSD at B) *A*=4.25 and C) *A*=9.52 are shown. The values of parameter *A* that generate these PSDs are denoted by vertical dotted lines in A) and D). The PSD of the model, in disparate regions of parameter space, captures the PSD of the alpha rhythm dynamics observed in the EEG data. D) Cost values (log scale) obtained from comparing the data and model PSDs, across the range of parameter *A*. This cost could only be calculated in regions of the model where a stable fixed point exists. Low cost values were obtained in regions on either side of the supercritical Hopf bifurcations.

When comparing model simulations to data, previous work has shown that even low-dimensional nonlinear models can give rise to non-convex cost landscapes that pose problems for model calibration (Kirk et al., 2008; Roesch & Stumpf, 2019). To illustrate this for NMMs, Fig. 1D quantifies the cost landscape of the model calibrated against eyes closed resting EEG for subject 1 (see Sec. 2.3 for a definition of the cost function used). This was calculated using grid sampling with *n*=1000 points across *A*=[0, 10], and through the linearised calculation of the PSD (see subsection 2.3). The figure shows that, as organised by the presence of bifurcations in the model, the cost landscape exhibits non-convexities, with regions of low cost occurring at disparate locations.

To further illustrate the non-convex cost landscapes present in the model, and begin to investigate the interplay of this landscape with bifurcations, we next tracked the supercritical Hopf and saddle-node bifurcations as parameters *A* and *a* are varied. Fig. 2A shows a bifurcation diagram as parameters *A* and *a* are varied. We note that a vertical cross-section at *a*=100 is equivalent to the bifurcation diagram shown in Fig. 1A. In Fig. 2A, the bifurcation diagram is shown superimposed on a cost heatmap across the 2 parameters varied, calculated using grid sampling with *n*=1000×1000 points across *A*=[0, 10] and *a*=[25,140]. The figure shows that the non-convexities illustrated previously persist in these 2 dimensions, with regions of low cost occurring on either side of the Hopf bifurcations (indicated by the red regions in the heatmap). Example PSDs generated from the model at 4 different parameter combinations are shown in Figs. 2B-E, highlighting that only the red regions of the cost landscape capture the same PSD shape seen in the EEG data. Additionally, Fig. S5 illustrates examples of cost landscapes across various 2-dimensional regions, highlighting the widespread occurrence of non-convexity in the model’s cost landscapes.

**Figure 2:**
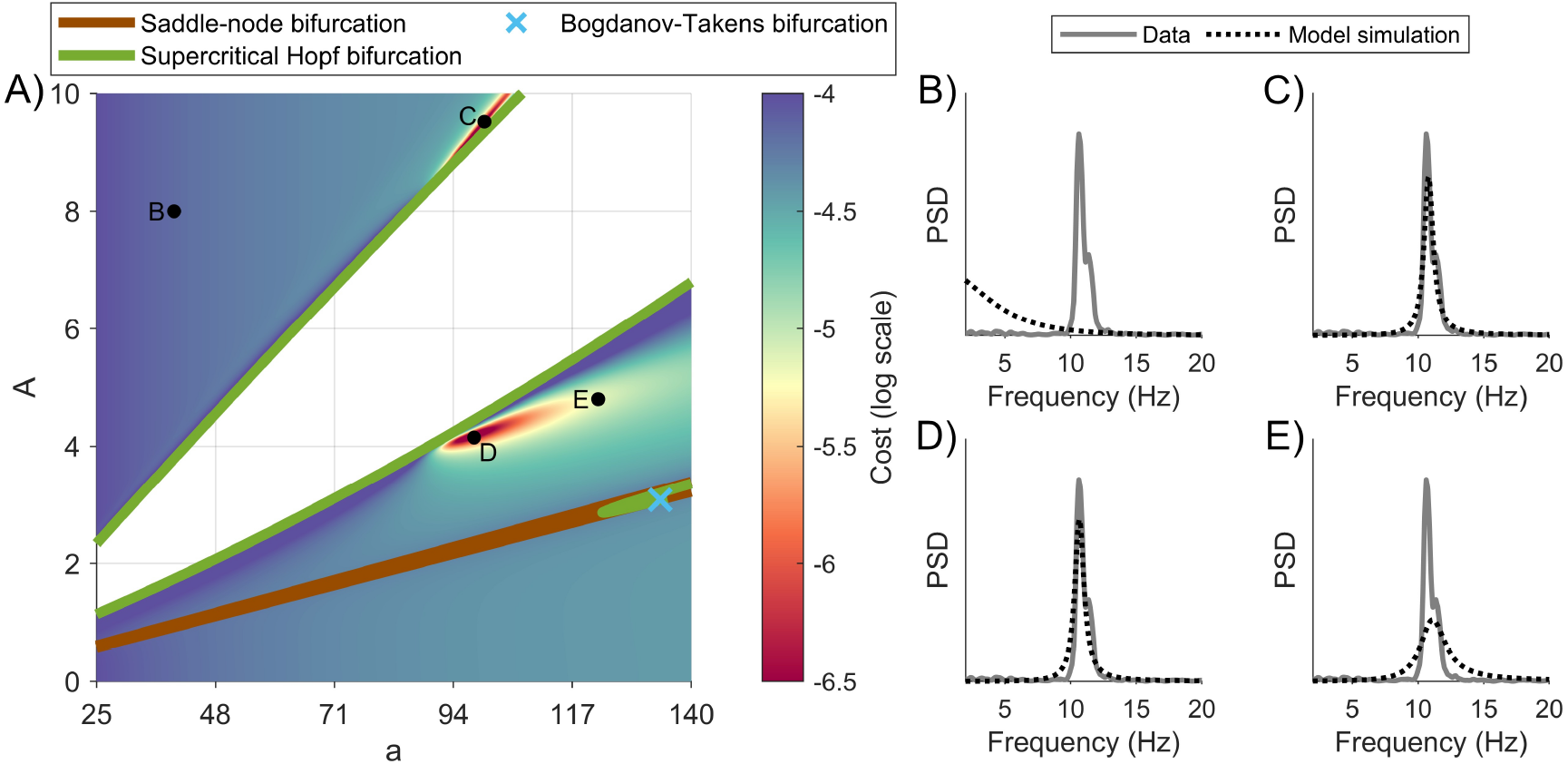
Bifurcation analysis and cost landscape of the Jansen and Rit model as parameters A and a are varied. A) The saddle-node and supercritical Hopf bifurcations are shown, along with a Bogdanov–Takens bifurcation (see legend). The bifurcation structure is overlaid on a 2-dimensional (log scale) cost heatmap, where red colours represent better fitting solutions between the model PSD (at the given parameter set) and the PSD of subject 1. We note that between the supercritical Hopf bifurcations (the region depicted in white), no stable fixed points were found, and hence the cost in this region could not be calculated from the linearised equations (see Sec. 2). The regions of low cost occur on either side of the supercritical Hopf bifurcations, indicating the non-convex cost landscape. Example model PSDs from the labelled points in A), superimposed with the EEG PSD from subject 1, are shown in B)-E). Other model parameters were set to their typical values, as defined in Table 1.

We have established that NMMs generate complex cost landscapes when their dynamics are compared to data. In the following section, we will evaluate algorithms to efficiently calibrate these models. Our focus is on identifying regions of parameter space that adequately reproduce the dynamics observed in EEG data. We quantify this by defining a threshold at which a model simulation is considered to pass visual inspection by being sufficiently similar to data (i.e. having a small enough cost value). Fig. 3A shows the EEG data PSD from subject 1, along with an example model simulation at the tolerance threshold defined for this subject. Simulations yielding a cost (see Equation 14) below this value are considered a good fit to the data. Fig. 3B shows the same cost heatmap presented in Fig. 2A, but with the set of LHS parameters that have been accepted for this subject (i.e. under the tolerance threshold) superimposed on the heatmap. Notably, the distribution of points under the threshold (which aligns with the cost landscape heatmap) would not be accurately modelled by a standard theoretical probability distribution. Additionally, Fig. 3C shows the LHS points recovered under the threshold when varying 3 model parameters (*A, a* and *b*). Similarly, the LHS points under the cost tolerance threshold exhibit a non-standard distribution. Figs. 3E-H show the same as Figs. 3A-D for a second example subject (subject 2). For this subject, the alpha rhythm observed is broader, and the location of the LHS points is consequently different. However, the distribution of LHS points under the cost tolerance threshold again does not conform to a standard probability distribution, as illustrated across 2 (Fig. 3E) and 3 (Fig. 3F) dimensions.

**Figure 3:**
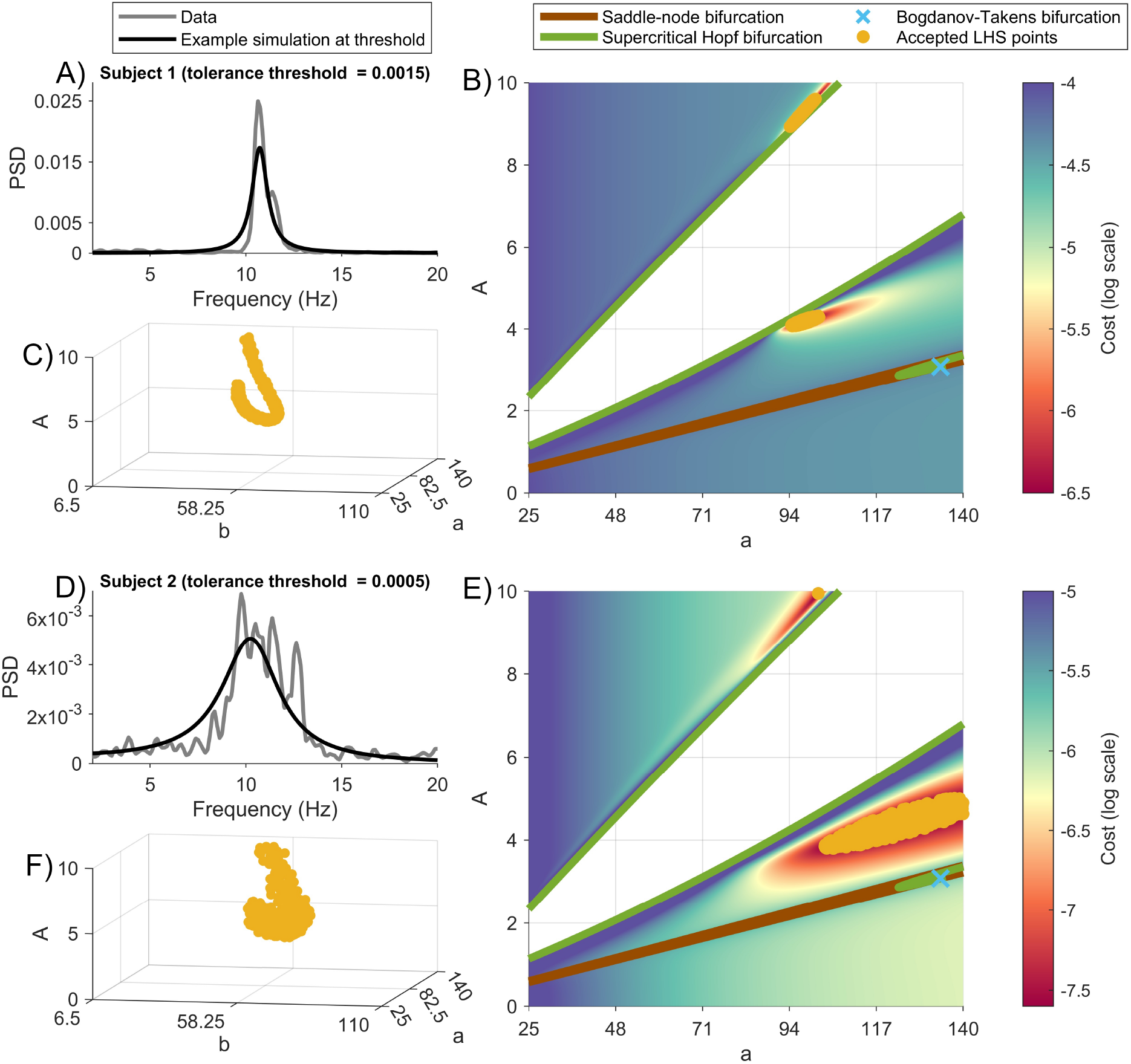
Examples of tolerance thresholds and accepted LHS points. A) PSD from data subject 1 and example model simulation at the tolerance threshold. This threshold quantifies an upper cost bound to a good fit of the model simulation to data. B) Cost heatmap and bifurcation structure of the model across parameters *A* and *a* (see Fig. 2 for details). LHS points that were accepted (i.e. parameter combinations that yielded simulations with a cost value under the defined threshold) are shown superimposed on the heatmap. C) Accepted LHS points across 3 parameters (*A, a* and *b*). In each case, other model parameters were set to their typical values, as defined in Table 1. For subject 2, D-F) shows the equivalent PSD, cost landscape, and accepted LHS points given in A-C), respectively.

### 3.2 Comparison of algorithms for model calibration from EEG data

Thus far, we have illustrated some of the complexities involved in calibrating NMM parameters from EEG data. Given that the distribution of parameters for which a model recreates data (within the defined threshold) would not be accurately modelled by a standard theoretical probability distribution, elucidating this set would clearly cause challenges for local search methods, or methods that rely on theoretical distributions. This therefore precludes the use of approaches such as DCM, which employs gradient descent-based optimisation and assumes Gaussian parameter distributions (i.e. the Laplace assumption). In what follows, we seek to compare three global algorithms for recovering regions of parameter space for which the model adequately simulates data. These global methods do not require a predefined theoretical probability distribution to be specified (see Sec. 2 for details). In addition, to ensure our results are not dependent on a particular subject or a particular tolerance threshold, we compare algorithms using EEG data from 10 subjects, and using four different tolerance threshold values.

The global algorithms we used were ABC, GA and NMMSO. We note that both ABC (Ditlevsen et al., 2023; West et al., 2021) and ESMs (Dunstan et al., 2023, 2025; Hartoyo et al., 2019, 2020; Nevado-Holgado et al., 2012; Wendling et al., 2005) have shown promise in recovering parameters of NMMs from M/EEG data. As detailed previously, we generated an estimate of the ground truth for comparing the algorithms by performing an LHS of the parameter space (see Sec. 2.5 for details). Moreover, we visually selected a tolerance threshold at which the dynamics of a model simulation recreate data. The comparison of the algorithms shown was conducted using the tolerance thresholds defined for each subject in Fig. S1. However, we also compared the algorithms using three different tolerance thresholds, with Figs. S6-S8 showing the parameter inference results obtained using each algorithm for tolerance thresholds defined in Figs. S2-S4, respectively.

Fig. 4 shows the parameters recovered from data using each of these algorithms, when fixing 7 of the 9 parameters in the model, and searching in a 2-dimensional region only (parameters *A* and *a*). In particular, Fig. 4A shows the points recovered using the LHS, ABC, GA and NMMSO methods to obtain parameter combinations that give rise to a model PSD that simulates the PSD observed in subject 1. The parameters obtained from each algorithm, depicted through a density approximation, and compared to the LHS, is shown in Fig. 4B. On visual inspection, for this subject, each algorithm appears to accurately recover regions of parameter space that can simulate the PSD of the data. We repeated this process on the EEG data of all 10 subjects. In Fig. 4C, we quantified the calibration accuracy by calculating the JSD for each subject, comparing the marginal density approximation of the recovered parameters from the LHS with the marginal density approximation of the recovered parameters from each algorithm tested (see Sec. 2.6 for details on the JSD). For both parameters *A* and *a*, we found that all algorithms had a relatively low JSD, with no significant differences between algorithms and a median JSD across subjects *<* 0.05 for every algorithm tested. Finally, Fig. 4D shows the number of model simulations (or function evaluations) executed using each method, with no significant differences observed between the algorithms. Thus, we conclude that the 3 methods perform equally well in 2 dimensions.

**Figure 4:**
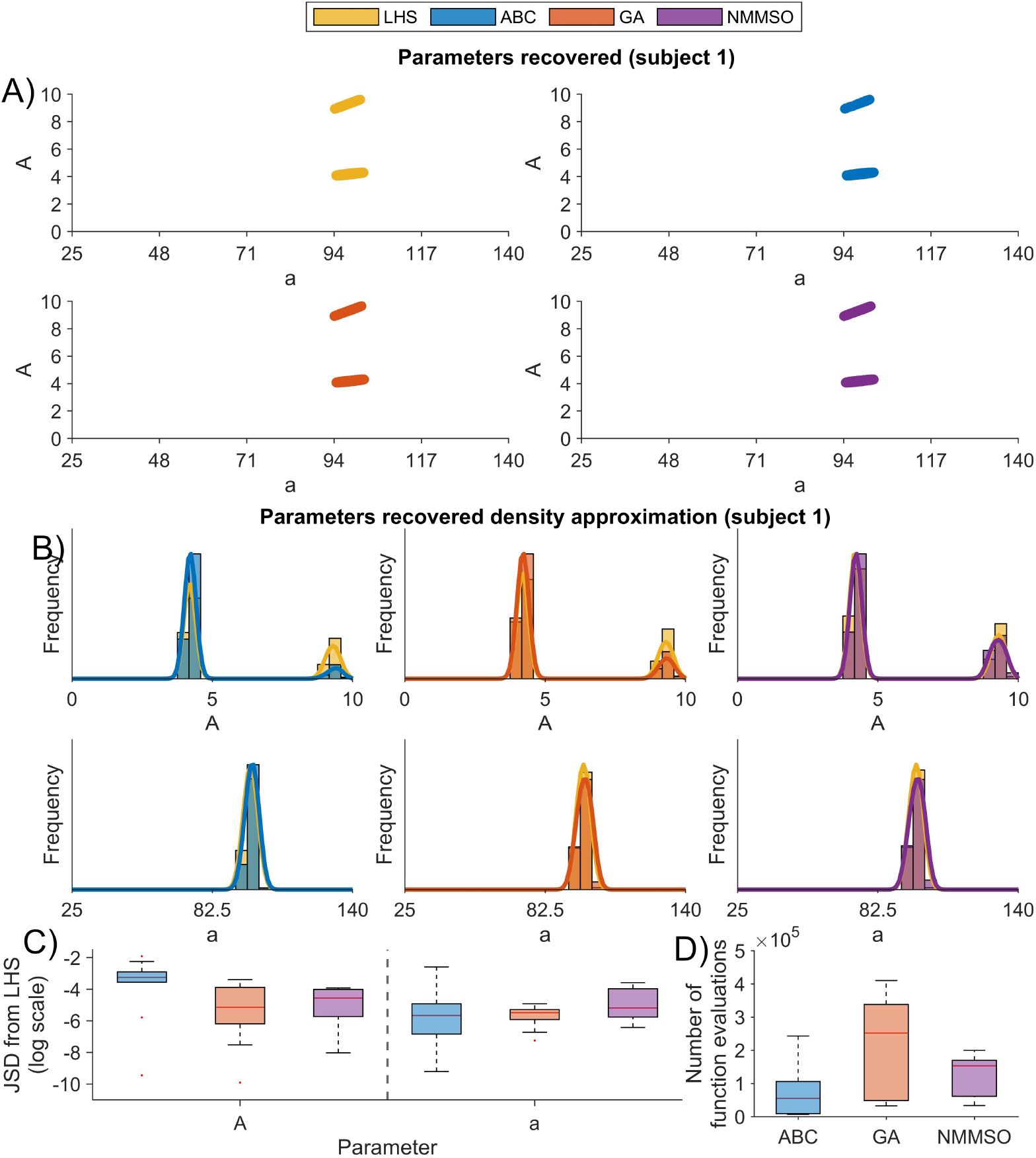
Comparison of algorithms for model calibration in 2 dimensions. A) Model parameters recovered from the LHS (top left), ABC (top right), GA (bottom left) and NMMSO (bottom right) for subject 1. Colours indicate regions where feasible parameters were found. B) Marginal density approximations obtained using each algorithm to recover parameters from the EEG data of subject 1. For each parameter, the x-axis limits are set to the parameter’s bounds. Each algorithm tested (ABC, GA and NMMSO) is shown superimposed on the approximation obtained from the LHS. In each case, the distributions consist of *N* =512 points and the model was calibrated by varying 2 parameters (*A* and *a*) within their bounded constraints, with other model parameters fixed to their typical values (see Table 1). C) JSD between the parameter distributions obtained from the LHS and the parameter distributions obtained from each method (log scale). D) The number of function evaluations used to execute each method. Note, no significant differences were observed in the Mann-Whitney U test after applying Bonferroni correction.

We next sought to compare the algorithms for model calibration in 3 dimensions. Fig. 5A shows the distribution of parameters recovered using each method on subject 1. Compared to the parameters recovered using ABC, the distributions recovered by the GA and NMMSO closely match those obtained through the LHS. In particular, ABC appears to only recover a subset of the parameters that describe the data, missing regions in the extremities. To visualise these distributions in more detail, Supplemental 2 shows a video of these points in 3 dimensions. Moreover, Fig. 5B shows the marginal density approximations obtained for this subject. We quantified differences between the algorithms across all subjects by calculating the JSD between the parameters recovered by the algorithm being tested and the parameters recovered from the LHS (Fig. 5C), along with the number of function evaluations used to execute each method (Fig. 5D). Here, no significant differences in the number of function evaluations were found, however, the GA and NMMSO had a significantly smaller JSD when compared to the LHS than the ABC’s JSD when compared to the LHS. This indicates that the parameter distributions obtained from the GA and NMMSO are more accurate than those obtained from the ABC when performing model calibration in 3 dimensions.

**Figure 5:**
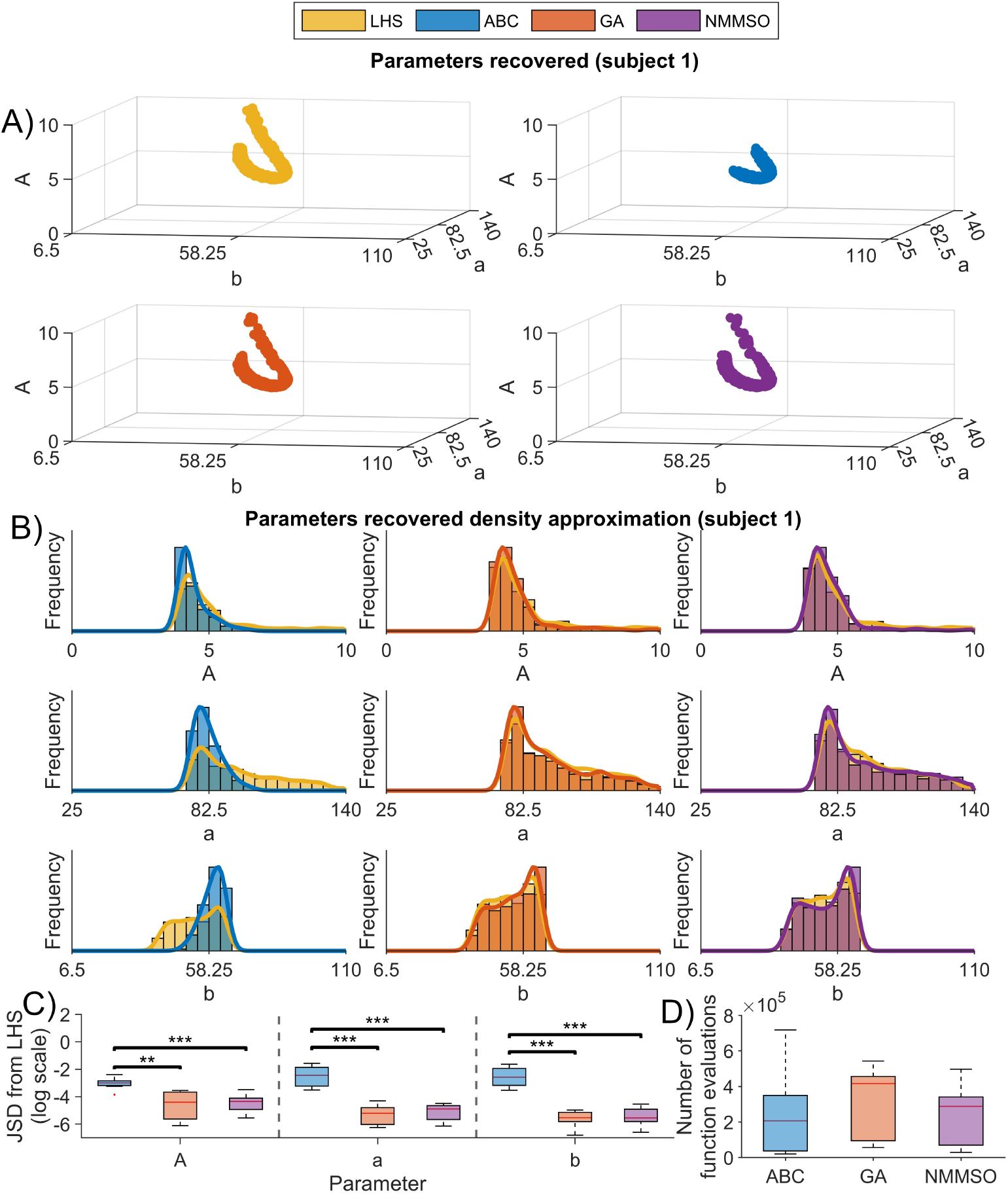
Comparison of algorithms for model calibration in 3 dimensions. A) Model parameters recovered from the LHS (top left), ABC (top right), GA (bottom left) and NMMSO (bottom right) for subject 1. Colours indicate regions where feasible parameters were found. B) Marginal density approximations obtained using each algorithm to recover parameters from subject 1. For each parameter, the x-axis limits are set to the parameter’s bounds. Each algorithm tested (ABC, GA and NMMSO) is shown superimposed on the approximation obtained from the LHS. In each case, the distributions consist of *N* =512 points and the model was calibrated by varying 3 parameters (*A, a* and *b*) within their bounded constraints, with other model parameters fixed to their typical values (see Table 1). C) JSD between the parameter distributions obtained from the LHS compared to the parameter distributions obtained from each method (log scale). D) The number of function evaluations used to execute each method. ^***^ *p <*0.001, ^**^ *p <*0.01 using a Mann-Whitney U test with Bonferroni correction.

In 3 dimensions, we have thus far found that the GA and NMMSO methods performed the most accurate model calibrations. Next, we sought to test how these algorithms compare when recovering parameters from the EEG data in the full 9 dimensions of the Jansen and Rit model. The marginal parameter distributions obtained when using the ABC, GA and NMMSO, in comparison to the LHS, are shown for subject 1 in Figs. S9-S11, respectively. Here, compared to ABC, it can be seen that the GA and NMMSO methods again perform better at recovering parameters that align with those obtained from the LHS (the “ground truth”). As before, we repeated this analysis across all subjects and quantified differences in the recovered parameter distributions by calculating the JSD between the parameters obtained from the LHS and each algorithm. The JSD values obtained across all subjects are shown in Fig. 6A. In general, the figure shows that the ESMs, and in particular the GA, obtained parameter distributions with the lowest JSD values compared to the LHS parameter distributions. Moreover, as shown by the function evaluations in Fig. 6B, the GA and NMMSO methods were able to perform the model calibration using fewer function evaluations than the ABC, with significance after Bonferroni correction only occurring for the comparison of the ABC and GA. Hence, the GA provided the most accurate model calibrations, while also being computationally efficient. We note here that these results were not dependent on the tolerance threshold used to define a good fit to data, with Figs. S6-S8 showing better performance of the GA compared to ABC when using a further three different tolerance threshold values.

**Figure 6:**
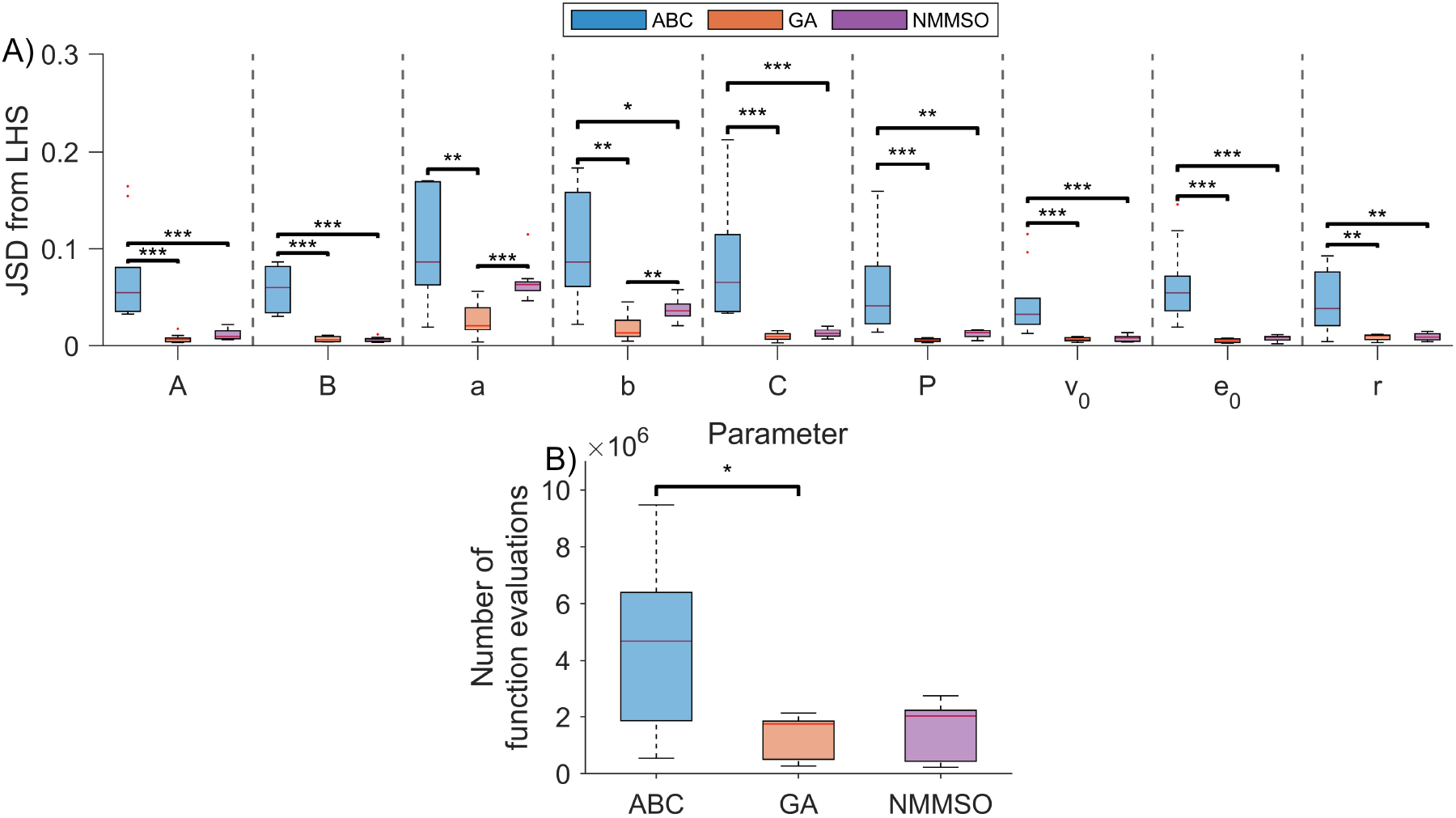
Comparison of algorithms for model calibration in 9 dimensions. A) JSD between parameter distributions recovered from the LHS (the “ground truth”) and parameter distributions recovered from each algorithm (see legend). B) The number of function evaluations used to execute each method. ^***^ *p <* 0.001, ^**^ *p <* 0.01, ^*^ *p <* 0.05 using a Mann-Whitney U test with Bonferroni correction.

### 3.3 Assessing parameter inference using a test cost function

Based on the analysis so far, we hypothesise that the poor inference obtained from ABC is due to the biased sampling introduced through iterating tolerance thresholds in the algorithm. For the cost landscapes presented, our results indicate that these biases make ABC more sensitive than the GA and NMMSO to suboptimal solutions. To test this, we designed a simple 1-dimensional cost function. This cost function can be seen in Fig. 7A. The function was designed such that 2 regions of the single design parameter, *x*, were below a cost tolerance threshold (similar to that seen for the NMM in Fig. 1). Moreover, the low cost region of *x* around *x*=0.95 contained a wider funnel to the minimum, whereas the low cost region of *x* around *x*=1.22 contained a narrower funnel to the minimum. We applied each parameter inference algorithm on this cost function in the same way as described previously. We refer to Sec. 2.7 for a definition of the cost function and the parameter input constraints. Our hypothesis was therefore that ABC could get stuck (i.e. wasted sampling) in regions that are locally good but globally suboptimal.

**Figure 7:**
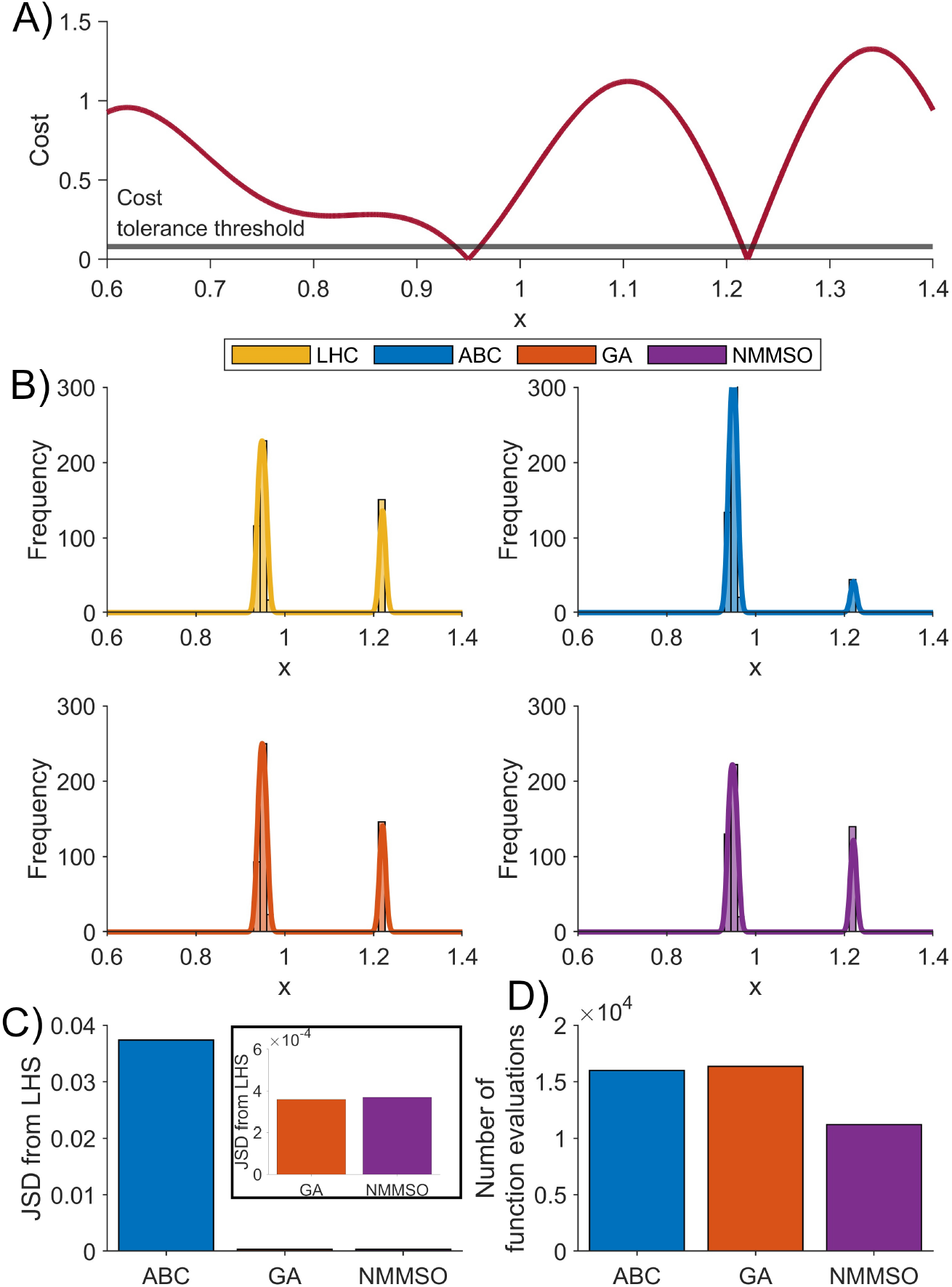
Comparison of algorithms for model calibration in a test cost function with a single parameter. A) 1-dimensional cost function. The function was designed such that 2 regions of the parameter space provide low cost (below the cost tolerance threshold line). The lower region contained relatively more nearby suboptimal solutions than the upper region. B) Model parameter distributions recovered from using the LHS (top left), ABC (top right), GA (bottom left) and NMMSO (bottom right) methods on the 1-dimensional test cost function. In each case, the distributions consist of *N* =512 points. C) JSD between the parameter distributions obtained from the LHS and the parameter distributions obtained from each method (see legend). The insert shows a different scale of the JSD to highlight the small values obtained from the GA and NMMSO methods. D) The number of function evaluations used to execute each method.

Fig. 7B shows the distribution of parameters obtained using each algorithm to recover *N* =512 points under the cost tolerance threshold. It can be seen that the GA (bottom left) and NMMSO (bottom right) methods generate distributions that visually resemble the distribution of parameter *x* found from using the LHS (top left). However, the distribution of *x* obtained from ABC (top right) appears to overestimate the relative density around *x*=0.95. In addition, in Fig. S12, we show the prior distributions from an implementation of the ABC across each iteration of the tolerance thresholds. Here, it can be seen that the larger funnel around *x*=0.95 causes a ‘biased’ prior around that area, which overestimates the required sampling in that region. These results therefore support our previous hypothesis regarding the poor inference obtained from ABC, and suggest that iterative tolerances commonly used in ABC to improve the algorithm’s acceptance rate make it sensitive to the shape of ‘funnels’ in the cost landscape. Our results suggest that both the GA and NMMSO are better designed to deal with complex cost landscapes, including these funnels. As before, these differences were quantified by the JSD (Fig. 7C) and the number of function evaluations executed for each algorithm (Fig. 7D), highlighting that the GA and NMMSO were the most accurate methods, with the NMMSO using the least number of function evaluations.

## 4 Discussion

In this study, we analysed the dynamical behaviour of the Jansen and Rit model and derived cost landscapes by comparing the model’s dynamics with empirical data. By quantifying cost in terms of the alignment of the model’s PSD with the PSD obtained from EEG data, we illustrated how parameter bifurcations organise the model’s dynamics and can give rise to non-convex cost landscapes that pose problems for model calibration. We found that regions of parameter space in which a model can accurately simulate data lie in the vicinity of supercritical Hopf bifurcations, which is consistent with the hypothesis that the brain operates near criticality (Cocchi et al., 2017). Furthermore, the non-convexities of the cost landscape mean that parameter combinations that can accurately simulate data would not be amenable to modelling using standard theoretical probability distributions. Our analysis therefore highlights the importance of implementing global methods when calibrating NMMs from EEG.

We subsequently compared the use of ABC and ESMs for calibrating the parameters of the Jansen and Rit model from EEG data. By comparing the regions of parameter space recovered from these algorithms to those obtained from high-resolution LHS (which represented a “ground truth”) across 2, 3 and 9 dimensions, we were able to evaluate the accuracy of each method. We quantified this by calculating the JSD between the parameter distributions recovered from each method and the parameter distributions obtained from the LHS. We observed that the ESMs, and in particular the GA, obtained distributions that had a lower JSD compared to the LHS distributions, than the distributions obtained from ABC. This indicates that the GA was the most accurate algorithm we tested for NMM calibration. Importantly, we found that despite these differences, the GA was still computationally efficient. In particular, when comparing the GA with the ABC, we observed no significant differences in the number of function evaluations in 2 and 3 dimensions, and a significant reduction in the number of function evaluations used to execute the GA in 9 dimensions.

### 4.1 Interpreting differences in the recovered parameter distributions

Bayesian inference methods have recently gained popularity in computational neuroscience (Baldy et al., 2024; Hadida et al., 2018; Jha et al., 2022). This includes ABC, which has been used to infer parameters of NMMs from data (Ditlevsen et al., 2023; West et al., 2021). ABC does not require the likelihood function to be analytically defined. This means ABC is particularly advantageous for models of brain activity, which are typically complex, nonlinear dynamical systems, whereby the evaluation of the likelihood function in a closed form is generally infeasible. Research on the use of ABC in the context of neural mass modelling has indicated that ABC is considerably affected by the curse of dimensionality (Jha et al., 2022). In line with this, we found that the inference obtained from ABC was substantially poorer in the 9-dimensional case, compared to the parameter inference performed in 2 and 3 dimensions. Here, we also investigated the mechanisms that explain this poor inference, highlighting that ABC is sensitive to the cost landscape (see Fig. 7 and Fig. S12).

An alternative method to ABC is to use ESMs—including the GA (Michalewicz, 1996) and NMMSO (Fieldsend, 2014) used herein. These algorithms are specifically designed for efficient exploration of high-dimensional spaces. They achieve this by iteratively refining candidate solutions through mechanisms such as genetic operations or swarm intelligence. The efficient search of the landscape requires balancing diversification, which helps to avoid being trapped in local optima, with intensification, which focuses on refining the current optimal solution estimates. By aligning the objective used in the ESMs with the defined summary statistic and distance measure in ABC, and observing when these values were minimised (within a defined tolerance threshold), we were able to compare these algorithms directly. We found that the multi-start method used in the ESMs could therefore enable the distributions recovered to be interpreted as approximations of probability distributions. We additionally showed that, compared to ABC, recovering parameters from data using a GA was more robust to the curse of dimensionality. In particular, for the GA, we observed only a subtle increase in the JSD from the distributions against the ground truth (the LHS) when varying 9 model parameters, compared to calibrating the model in the lower-dimensional cases. In addition to issues with high-dimensional inference, it has been shown that ABC algorithms are susceptible to inaccurate inference when the true posterior lies far from the prior (Lintusaari et al., 2016). For the application of neural mass modelling of EEG, there is often little information available that helps us to select informed priors on many of the parameter values (Goodfellow et al., 2022). This is largely because of the issues of measuring parameters at the scale of mean activity from neural populations of brain tissue. Hence, herein we used uninformative prior information to initialise the algorithms tested. For the ESMs, this involved sampling from large regions of parameter space within box constraints. In the case of ABC, this involved using uniform priors between the parameter bounds. It is therefore likely that a large deviation from the prior is needed to obtain the true posterior, and hence our results are consistent with previous findings (Lintusaari et al., 2016). However, we additionally propose that GAs are a useful alternative in scenarios where there is no (or little) prior information on parameters available. Since Bayesian algorithms provide an efficient means of incorporating prior information, it would be interesting in the future to compare calibration methods for NMM parameters in scenarios where more prior information is available. Moreover, future work will probe the use of dynamics-informed priors for Bayesian inference, whereby ESMs are used to help construct the priors for Bayesian algorithms.

### 4.2 Assumptions and limitations

In this study, we made three key assumptions when performing model calibration on NMMs. These were: (i) focusing on the Jansen and Rit model, (ii) linearising the model and thereby focusing attention on dynamics close to fixed points, and (iii) comparing model dynamics to data solely in terms of a comparison of the PSD. These assumptions were important to enable a tractable and feasible approach for comparing the algorithms we tested. We used the Jansen and Rit model as it is an archetypal NMM. We note that many of the dynamical properties observed in the Jansen and Rit model will generalise to other NMMs (Cooray et al., 2023). However, despite having 9 parameters, the Jansen and Rit model is relatively low-dimensional, so it would be interesting to test how the algorithms studied herein scale for NMMs with more parameters and a greater repertoire of dynamical behaviour. Furthermore, although the brain is largely believed to operate in a linear regime during a resting state (Stam et al., 1999), this is not exclusively the case, and research has recovered nonlinear dynamics from resting data (Dunstan et al., 2023). Moreover, other states such as seizures are nonlinear phenomena (Lehnertz, 2008). In general, it is therefore important to consider nonlinear measures, both independently and alongside the PSD analysed in this study, when comparing model dynamics to data. One benefit of evolutionary computation is that a well-established framework exists for using these algorithms for multi-objective optimisation. In the future, methods for recovering parameters of NMMs from brain imaging data should be contrasted by evaluating multiple objectives between model simulations and data (Dunstan et al., 2023, 2025).

### 4.3 Conclusion

By analysing dynamical properties of an archetypal NMM, and quantifying cost landscapes, this study illustrated some of the complexities involved in calibrating models using brain imaging data. We found that model parameter sets that give rise to dynamics seen in EEG data would not be accurately modelled by standard theoretical probability distributions. We therefore evaluated the performance of alternative global model calibration methods that do not assume fixed form posteriors. Our findings indicate that ESMs are useful tools for dealing with the complex cost landscapes that arise, and for providing accurate NMM parameter calibration.

## Data and Code Availability

Code supporting the findings is publicly available and maintained as a GitHub repository (https://github.com/domdunstan/Global_NMM_parameter_inference). The full EEG dataset is publicly available (https://osf.io/f2vya), with the processed power spectral data available within the GitHub repository.

## Ethics

All participants gave written informed consent before enrolment.

## Author Contributions

**DMD**: Conceptualization, Methodology, Software, Formal analysis, Writing - Original Draft, Writing - Review & Editing, Funding acquisition. **MPR**: Investigation, Writing - Review & Editing **JEF**: Conceptualization, Software, Writing - Review & Editing, Supervision. **MG**: Conceptualization, Methodology, Writing - Review & Editing, Supervision, Funding acquisition.

## Funding

DMD acknowledges the generous financial support provided via an Early Career Researcher Funding project under EPSRC grant EP/T017856/1 and an EPSRC DTP studentship (ref 2407565). Data collection was funded by Medical Research Council (MRC) Programme grant number MR/K013998/1.

## Licence

For the purpose of open access, the author has applied a Creative Commons Attribution (CC BY) licence to any Author Accepted Manuscript version arising from this submission.

## Declaration of Competing Interests

The authors declare no conflict of interest.

## Acknowledgements

We would like to thank Margaritis Voliotis for providing helpful comments on the manuscript.

## Supplementary Material

See “Supplemental.pdf” for supplemental figures and “Supplemental 2.mp4” for a supplemental video.

## Notes

### Competing Interest Statement

The authors have declared no competing interest.

