## Supplemental for "Global search metaheuristics for neural mass model calibration"

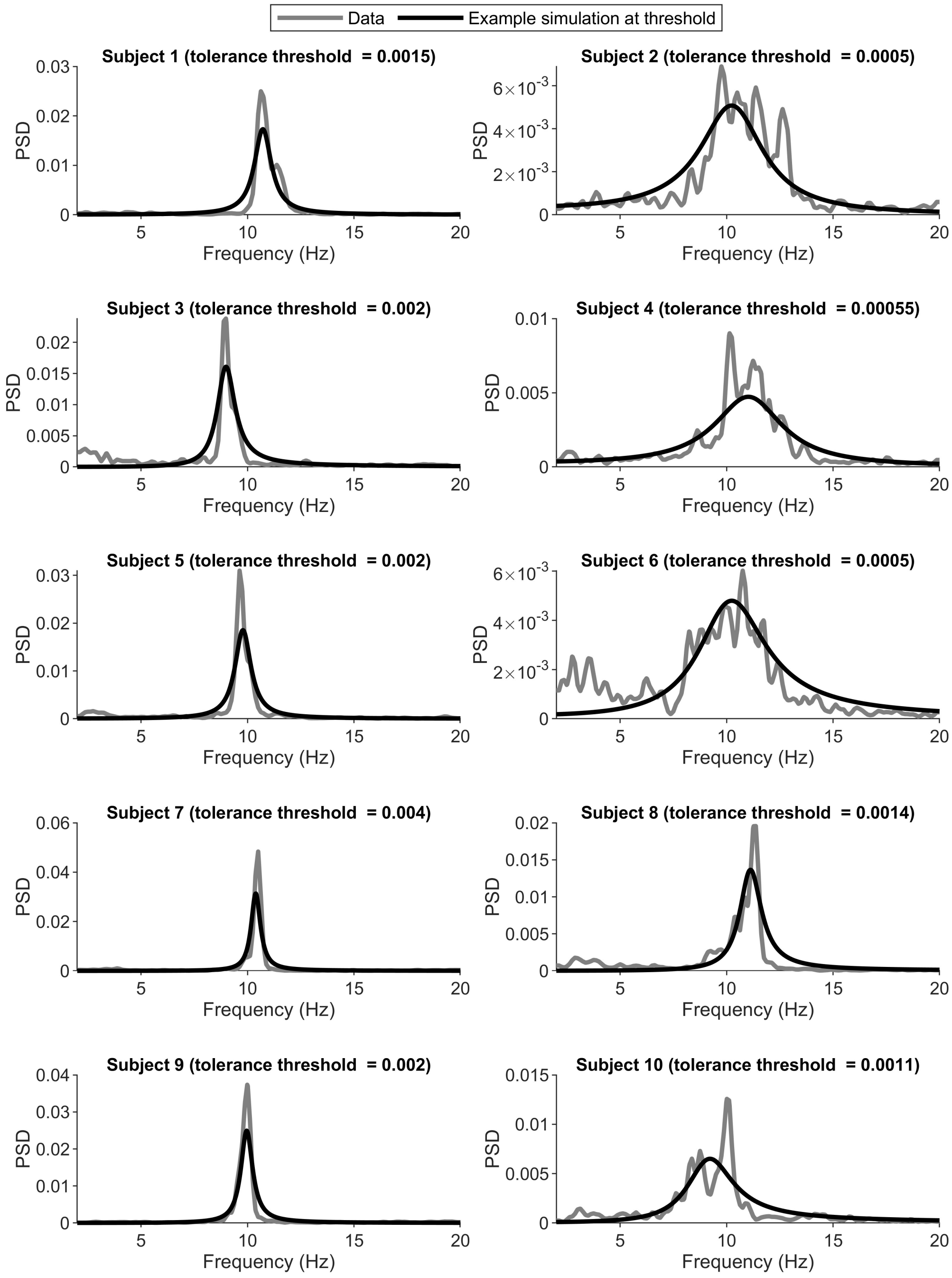


Figure S1: **Data PSD and example model PSD at cost tolerance threshold.** Each subplot shows a different subject. The model PSDs show an example simulation at the cusp of the tolerance threshold (as defined for each subject in the corresponding subplot title). Simulations under this cost are regarded as a sufficiently good fit to the data.

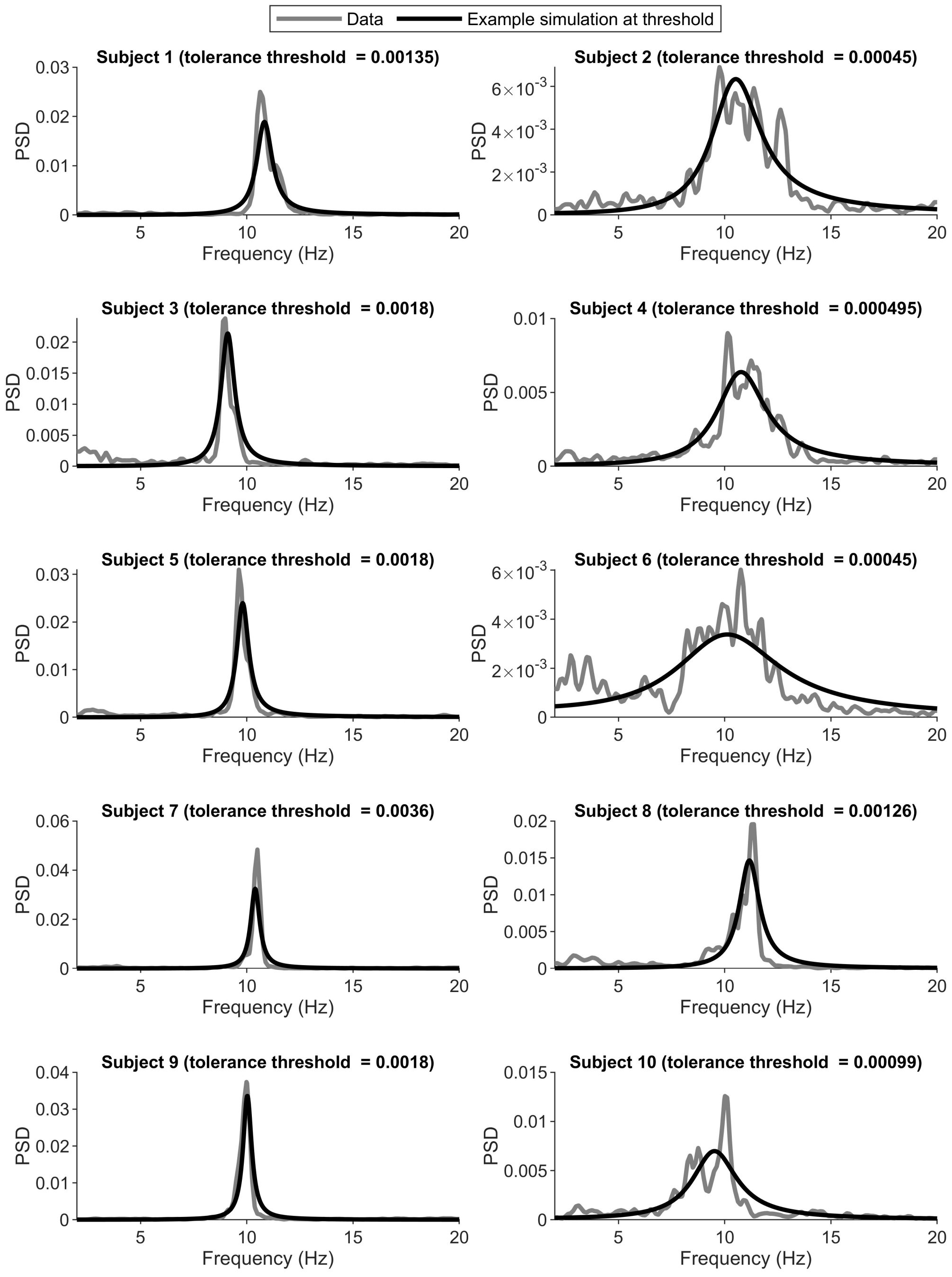


Figure S2: **Data PSD and example model PSD at cost tolerance threshold 2.** Each subplot shows a different subject. The model PSDs show an example simulation at the cusp of the tolerance threshold (as defined for each subject in the corresponding subplot title). Simulations under this cost are regarded as a sufficiently good fit to the data. This example shows a tolerance 10% stricter than Fig. S1.

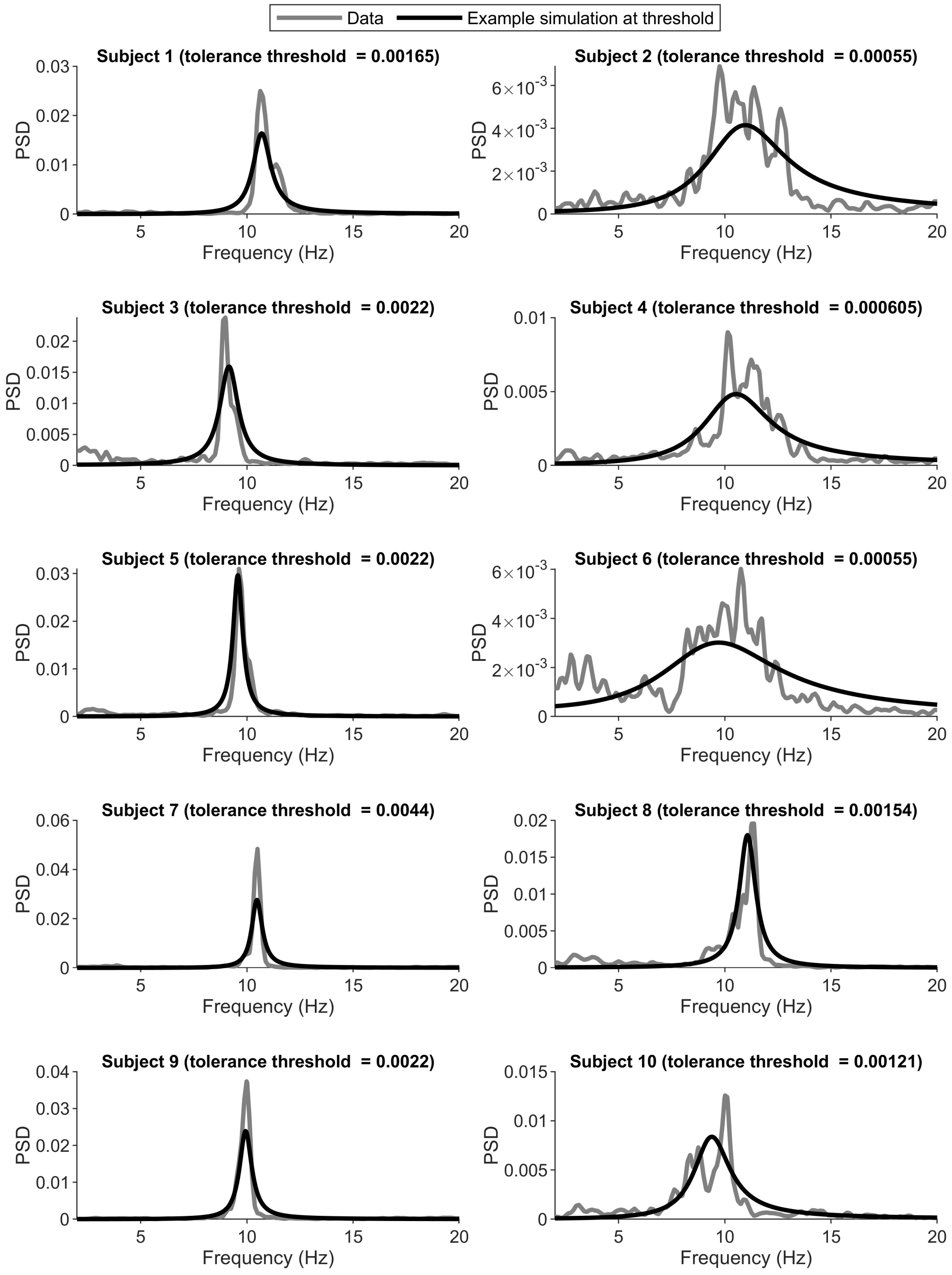


Figure S3: **Data PSD and example model PSD at cost tolerance threshold 3.** Each subplot shows a different subject. The model PSDs show an example simulation at the cusp of the tolerance threshold (as defined for each subject in the corresponding subplot title). Simulations under this cost are regarded as a sufficiently good fit to the data. This example shows a tolerance 10% more lenient than Fig. S1.

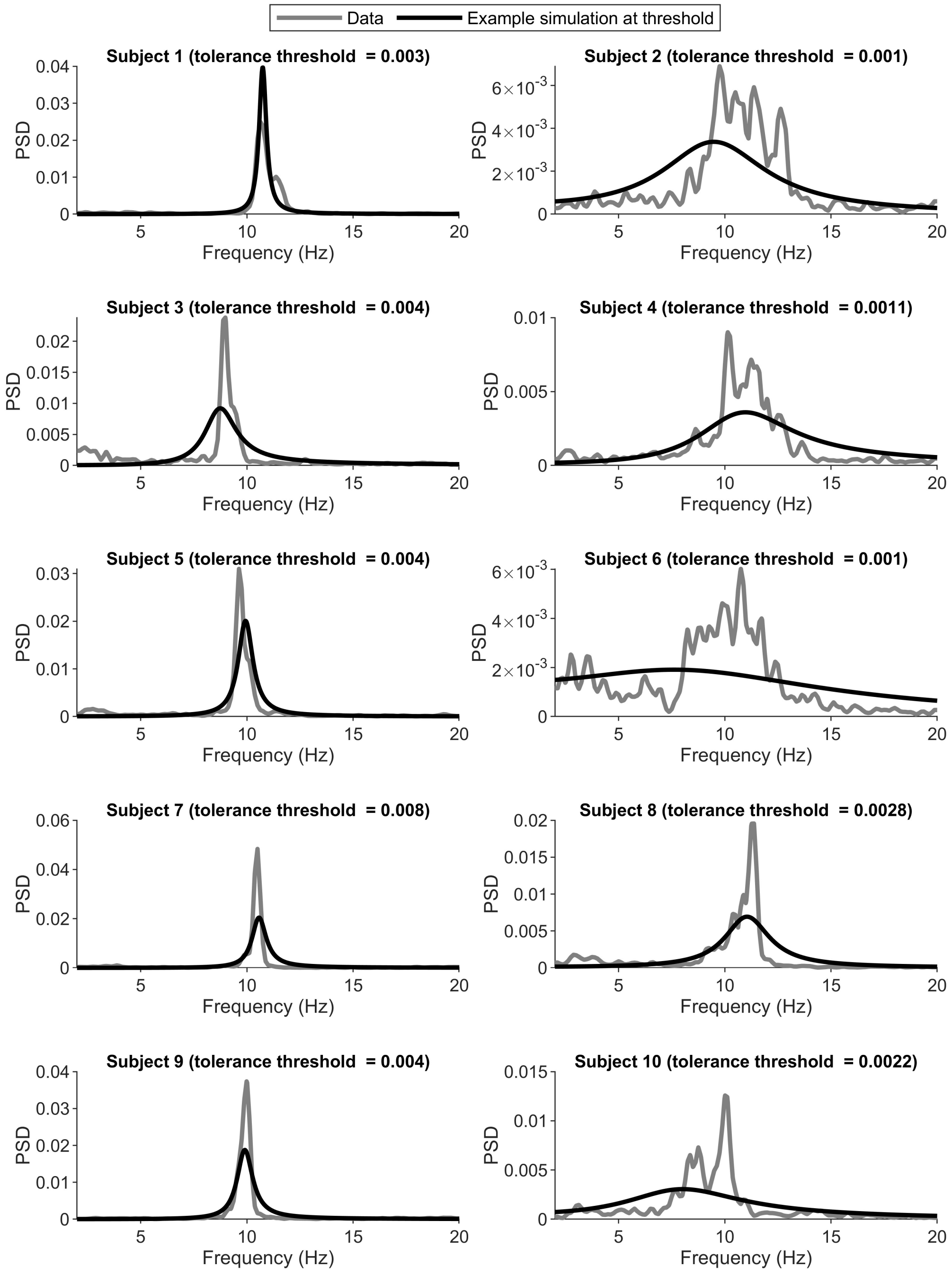


Figure S4: **Data PSD and example model PSD at cost tolerance threshold 4.** Each subplot shows a different subject. The model PSDs show an example simulation at the cusp of the tolerance threshold (as defined for each subject in the corresponding subplot title). Simulations under this cost are regarded as a sufficiently good fit to the data. This example shows a tolerance 100% more lenient than Fig. S1.

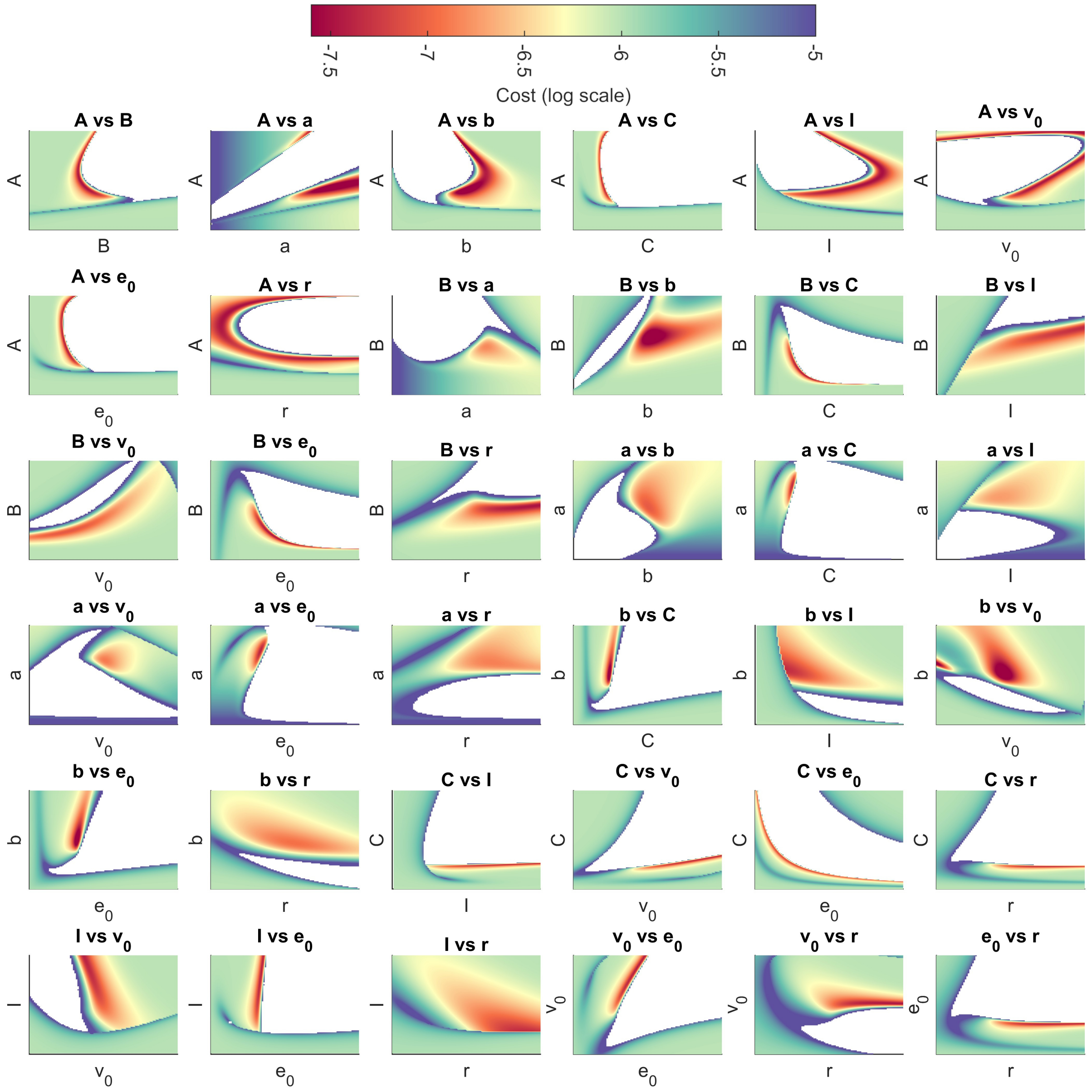


Figure S5: **Cost landscape of the Jansen and Rit model across pairwise model parameter combinations.** Subplot titles (and axes) denote the parameter combinations, with other model parameters set to their typical values (see Table 1 in the Manuscript). Axis limits are set to the bounds defined in Table 1 of the Manuscript. The cost (on a log scale and measured by the sum of squared error) shows how well the model at the given parameter combination simulates the data PSD. These figures were generated from comparing model simulations to the PSD estimated from the EEG of subject 2. The white region shows an area where no stable fixed points were found, and hence the cost could not be calculated from the linearised equations in this region.

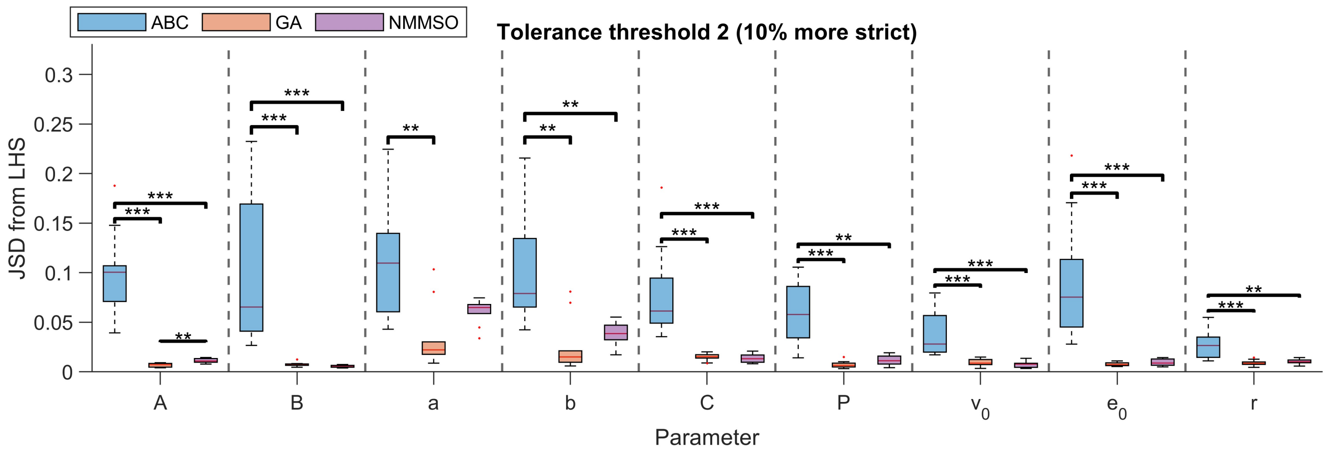


Figure S6: **JSD from tolerance thresholds defined in Fig. S2.** The JSD between parameter distributions obtained from an LHS and each algorithm is shown (see legend). These were obtained from varying all 9 parameters in the Jansen and Rit model and using the tolerance threshold defined in Fig. S2 (10% stricter). ***$p < 0.001$, **$p < 0.01$, *$p < 0.05$ using a Mann-Whitney U test with Bonferroni correction.

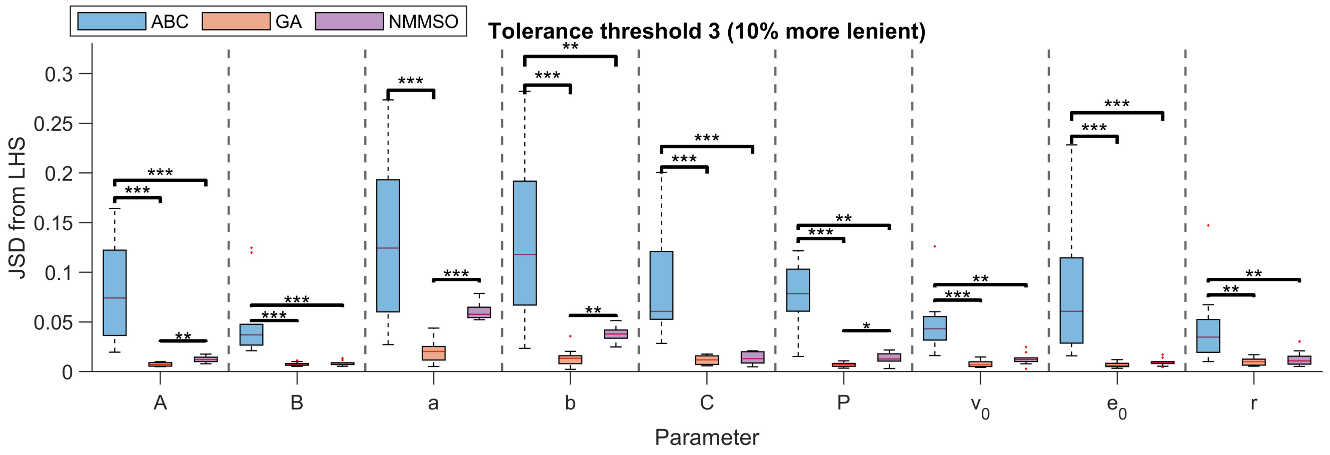


Figure S7: **JSD from tolerance thresholds defined in Fig. S3.** The JSD between parameter distributions obtained from an LHS and each algorithm is shown (see legend). These were obtained from varying all 9 parameters in the Jansen and Rit model and using the tolerance threshold defined in Fig. S3 (10% more lenient). ***$p < 0.001$, **$p < 0.01$, *$p < 0.05$ using a Mann-Whitney U test with Bonferroni correction.

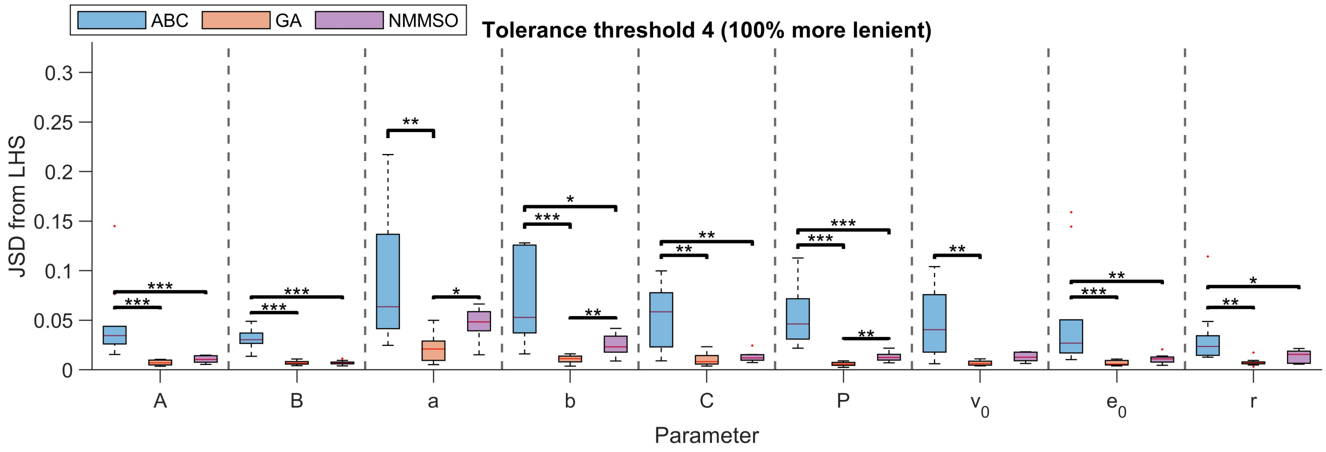


Figure S8: **JSD from tolerance thresholds defined in Fig. S4.** The JSD between parameter distributions obtained from an LHS and each algorithm is shown (see legend). These were obtained from varying all 9 parameters in the Jansen and Rit model and using the tolerance threshold defined in Fig. S4 (100% more lenient). ***$p < 0.001$, **$p < 0.01$, *$p < 0.05$ using a Mann-Whitney U test with Bonferroni correction.

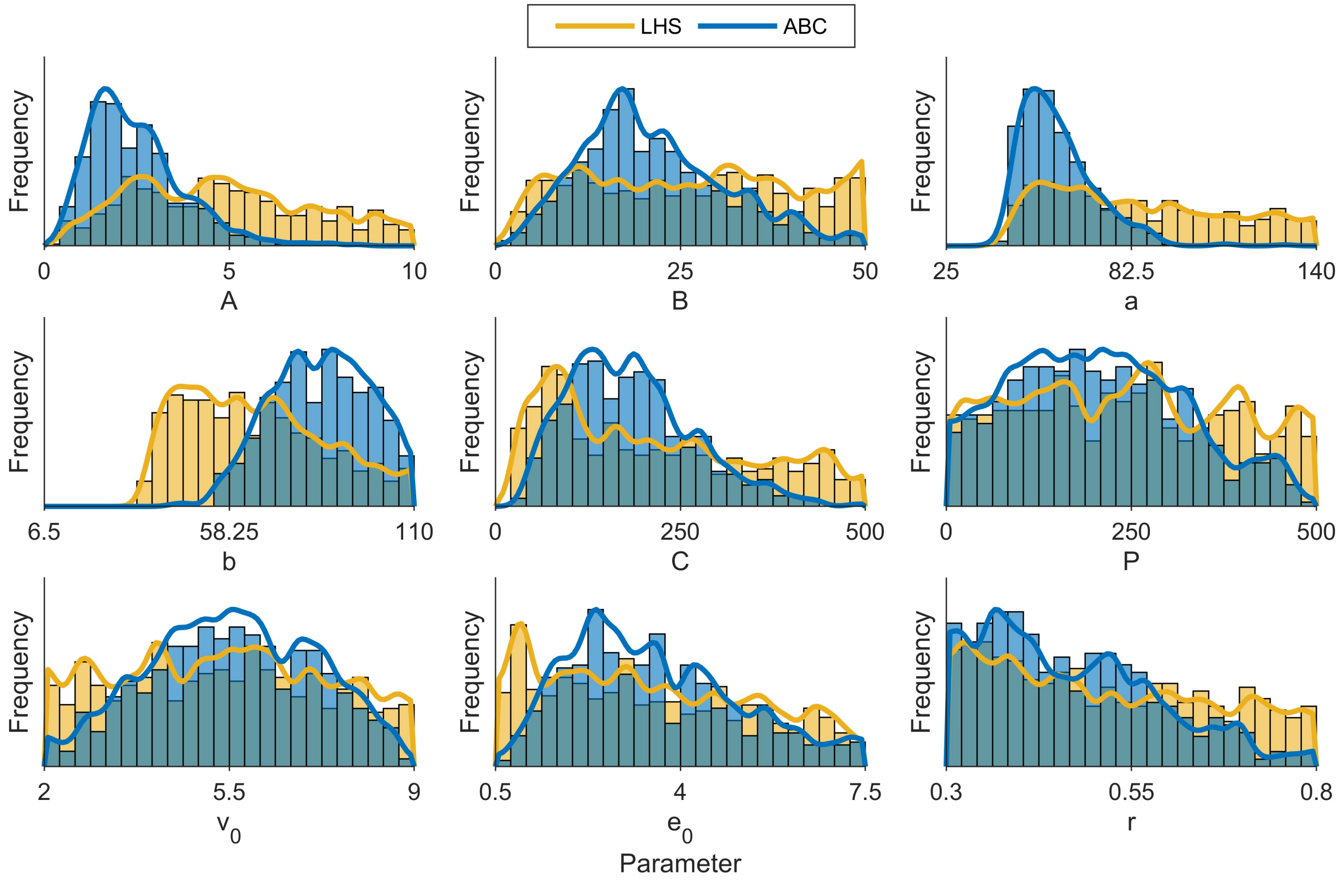


Figure S9: **Marginal parameter density approximation for subject 1, obtained using ABC in 9-dimensions.** The distributions obtained from ABC are shown superimposed on the parameter distributions obtained from LHS (see legend). For each parameter, the x-axis limits are set to the parameter's bounds.

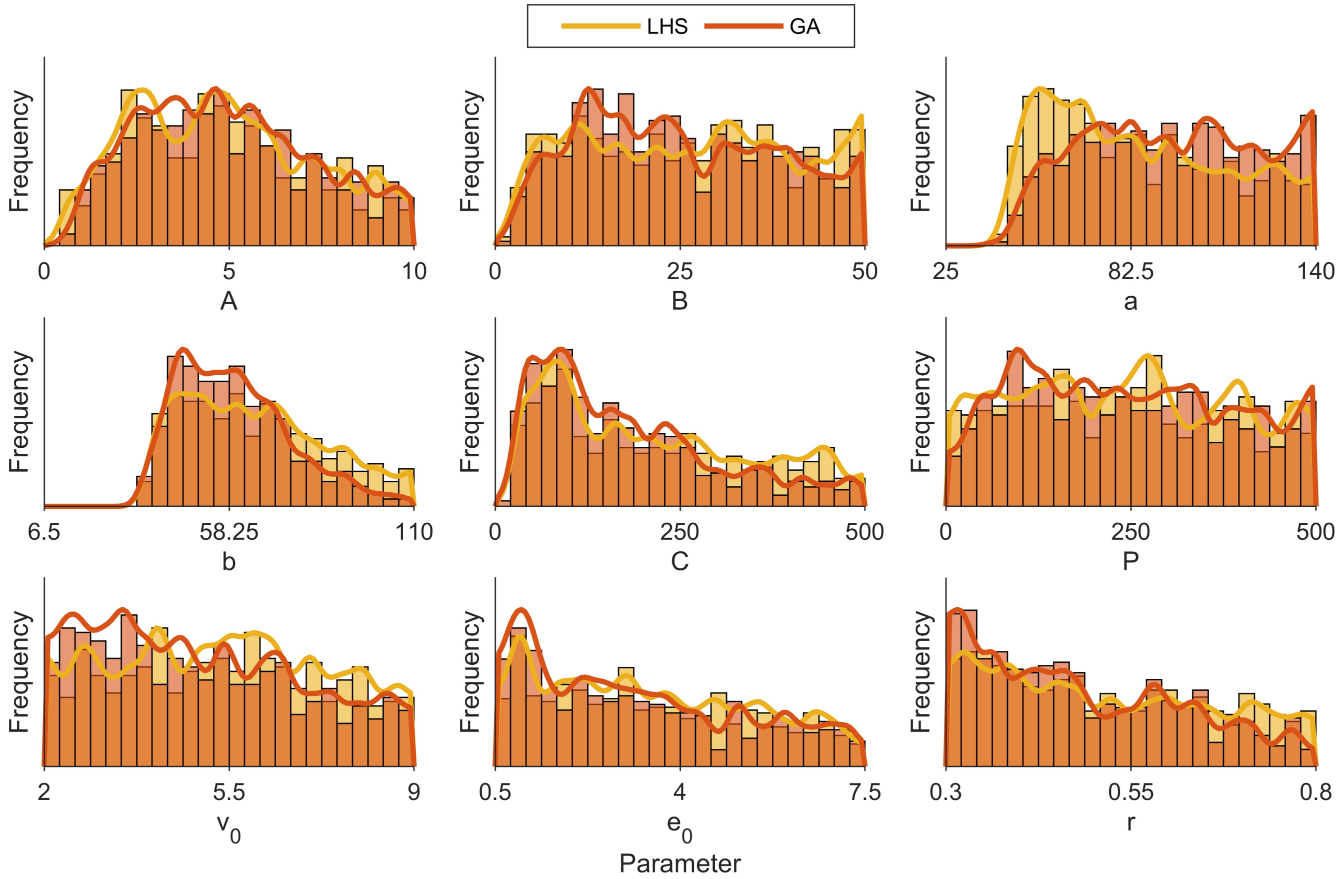


Figure S10: **Marginal parameter density approximation for subject 1, obtained using GA in 9-dimensions.** The distributions obtained from GA are shown superimposed on the parameter distributions obtained from LHS (see legend). For each parameter, the x-axis limits are set to the parameter's bounds.

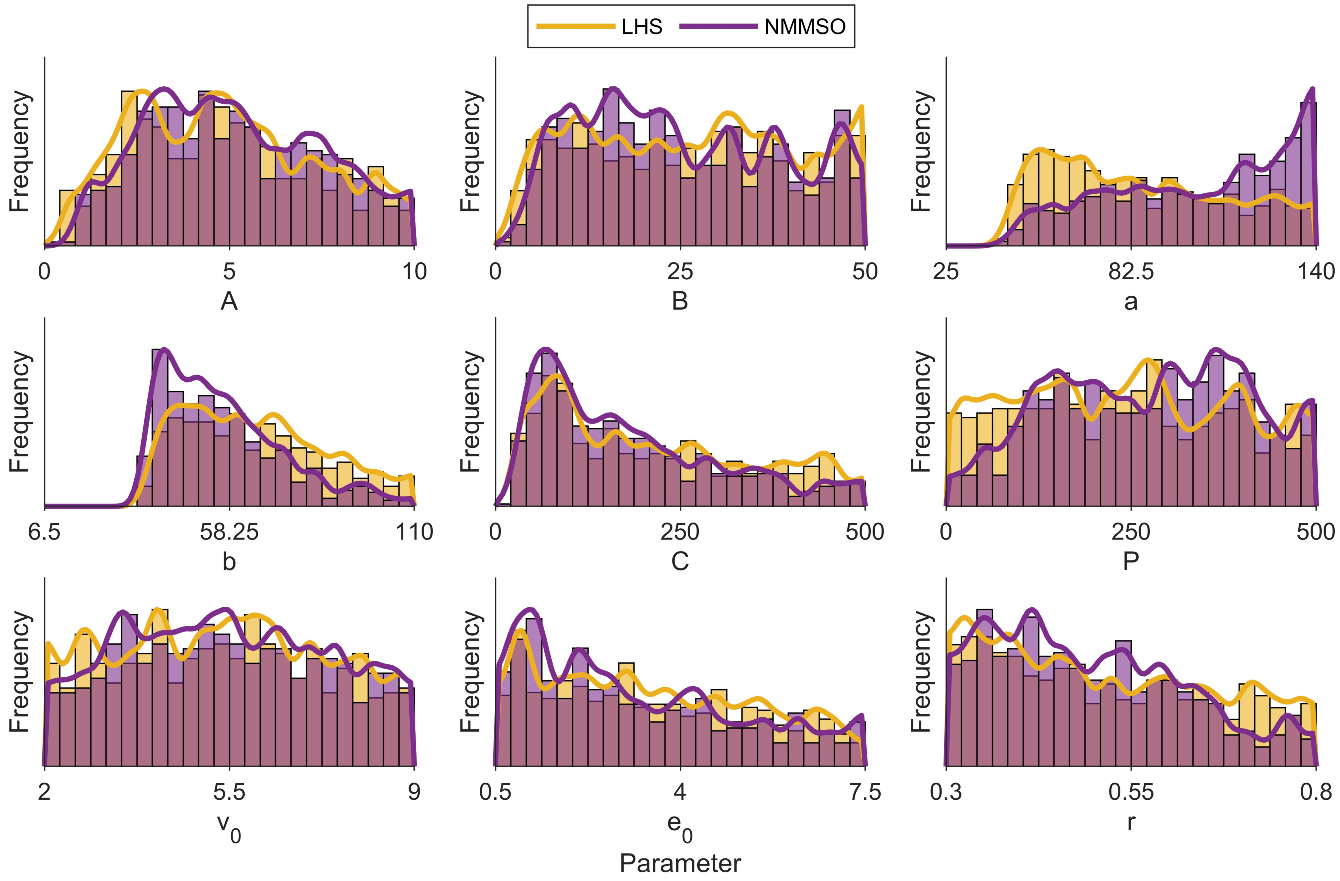


Figure S11: **Marginal parameter density approximation for subject 1, obtained using NMMSO in 9-dimensions.** The distributions obtained from NMMSO are shown superimposed on the parameter distributions obtained from LHS (see legend). For each parameter, the x-axis limits are set to the parameter's bounds.

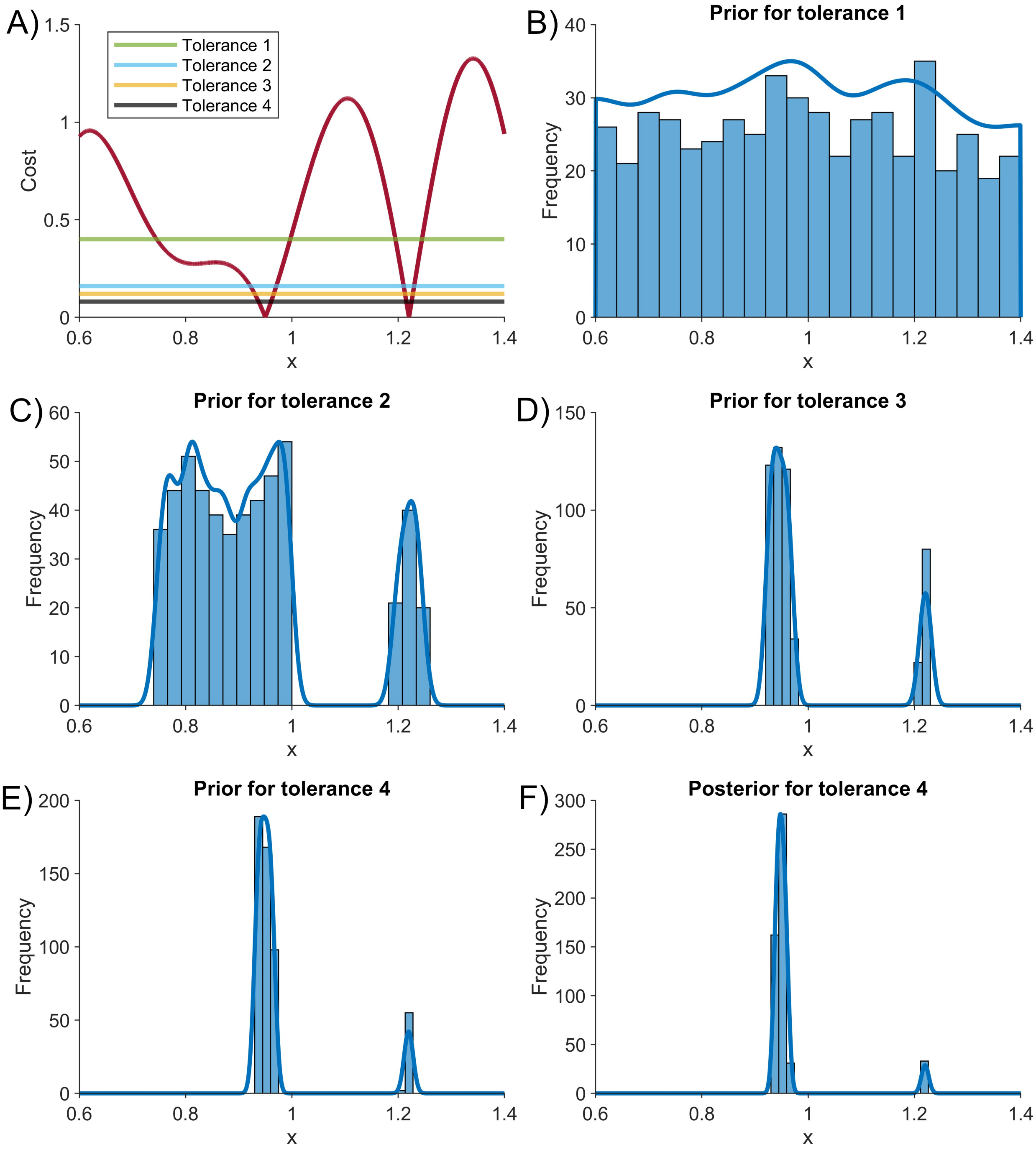


Figure S12: **Tolerance schedule of ABC on one-dimensional cost function.** A) One-dimensional cost function (see Methods in Manuscript for definition). A horizontal line representing the cost for each tolerance used in the schedule has been added. Tolerance 4 represents the final cost tolerance threshold value. B-E) Iterative priors used for each of the cost tolerances (see subtitles). Note, the first prior involves taking draws from a uniform distribution between the box constraints. F) Posterior from the final tolerance. This posterior deviates substantially from that obtained using the LHS (see Manuscript). The ABC method used does not provide a robust inference due to overestimating density around the lower region of low cost, as a consequence of biasing the priors around this region.
